# An unconventional interaction interface between the peroxisomal targeting factor Pex5 and Eci1 enables PTS1 independent import

**DOI:** 10.1101/2024.06.02.597006

**Authors:** Lior Peer, Nadav Elad, Orly Dym, Asa Tirosh, Jossef Jacobovitch, Shira Albeck, Maya Schuldiner, Yoav Peleg, Einat Zalckvar

## Abstract

Accurate and regulated protein targeting to organelles is crucial for eukaryotic cellular function and homeostasis. This has driven the evolution of targeting signals on proteins and the targeting factors that recognize them. One example for this is peroxisomal matrix proteins, the majority of which rely on the targeting factor Pex5 to correctly localize and function. While most Pex5 cargos contain a Peroxisomal Targeting Signal type 1 (PTS1), in recent years it has become clear that more binding interfaces exist, and that targeting by Pex5 is more complex than previously thought. Here, we uncover that the matrix protein Eci1 can reach peroxisomes in the absence of its PTS1. By solving the structure of a complex between full length yeast Pex5 and Eci1 using Cryo-Electron Microscopy, we could identify their binding interfaces. This allowed us to map an additional binding interface that is independent of the canonical PTS1-mediated binding site. Our work brings forward a solution to a long-standing mystery regarding Eci1 targeting to peroxisomes. More globally, it demonstrates the intricate and complex nature of organelle targeting and how it has evolved to serve the complex eukaryotic environment.

## Introduction

Cells produce a plethora of proteins, many of which must be localized into specific cellular compartments to function properly and enable cellular homeostasis (Aviram and Schuldiner, 2017; Bykov *et al*., 2020). When a protein is mistargeted, it can lead to severe cellular implications: First, the protein is absent from its target compartment, leading to loss of function; Second, the protein can be mislocalized to an alternate compartment, causing a toxic gain of function; Finally, the protein may aggregate, causing cellular stress and cytotoxicity. To avoid these deleterious effects, cells have evolved intricate pathways to ensure correct spatial and temporal protein targeting.

One cellular compartment whose proper function relies on correct protein localization is the peroxisome. Peroxisomes are vital organelles, carrying important metabolic functions and processes such as the catabolism of fatty acids, detoxification of reactive oxygen species (ROS) and synthesis of plasmalogen (Islinger *et al*., 2018). Aberrant peroxisomal function or lack of peroxisomes results in severe diseases, such as Zellweger spectrum disorders (Argyriou, D’Agostino and Braverman, 2016; Waterham, Ferdinandusse and Wanders, 2016). Moreover, in recent years, it has become clear that deterioration in peroxisomal function is linked to prevalent pathological conditions such as diabetes, neurodegeneration, cancer, and others (Zalckvar and Schuldiner, 2022; Wanders *et al*., 2023).

Since peroxisomes do not possess their own genome, all peroxisomal matrix (lumen) proteins are nuclear encoded, synthesized on cytosolic ribosomes and must be targeted and imported. To properly target, peroxisomal matrix proteins often contain a canonical Peroxisomal Targeting Signal Type 1 (PTS1, located at the C terminus) or Type 2 (PTS2, located close to the N terminus). The peroxisomal cargo factors Pex5 and Pex9 identify the targeting signal PTS1 (Effelsberg *et al*., 2016; Yifrach *et al*., 2016), while Pex7 recognizes PTS2 (Walter and Erdmann, 2019).

While most Pex5 cargo has been shown to have a PTS1, it is now becoming clear that numerous Pex5 cargo proteins do not have a PTS1 (Klein *et al*., 2002; Kempiński *et al*., 2020; Yifrach *et al*., 2022). Moreover, some PTS1 proteins can be targeted when their PTS1 is masked (i.e., not exposed in the cytosol and/or unavailable for Pex5 recognition) (Kempiński *et al*., 2020; Yifrach *et al*., 2022). These observations indicate that Pex5 can target proteins in a PTS1 independent manner.

One such protein that Pex5 targets in a PTS1-independent manner is the yeast Eci1 (delta3, delta2-Enoyl-CoA Isomerase 1), an enzyme that is essential for the beta-oxidation of unsaturated fatty acids in peroxisomes. While Eci1 has a PTS1 (Geisbrecht *et al*., 1998; Gurvitz *et al*., 1998), it is clear that it does not require it for targeting. The PTS1 independent targeting mechanism of Eci1 has therefore become a source of debate (Geisbrecht *et al*., 1999; Karpichev and Small, 2000; Yang, Purdue and Lazarow, 2001). Two hypotheses for the enigmatic targeting mechanism of Eci1 have been suggested: The first is that Eci1 has both a PTS1 and a PTS2 (Karpichev and Small, 2000; Yang, Purdue and Lazarow, 2001). The second hypothesis suggests that when its own PTS1 is masked or absent, Eci1 can “piggyback” on its paralog, Dci1 (Geisbrecht *et al*., 1999; Karpichev and Small, 2000; Yang, Purdue and Lazarow, 2001). Nevertheless, neither hypothesis were unequivocally proven and the mechanisms by which Eci1 is targeted to peroxisomes in the absence of its own PTS1 remained a mystery.

Here, we solve this long-standing conundrum. We show that indeed Eci1-monomeric NeonGreen (mNG) is localized to peroxisomes despite having its PTS1 masked by the mNG fluorophore. We demonstrated that under these circumstances, Eci1 still uses only Pex5 and that its targeting cannot be fully accounted for by piggybacking on Dci1. These observations supported a third hypothesis proposing the presence of an additional non-canonical binding interface between Pex5 and Eci1. To test this hypothesis, we co-expressed the full-length yeast Pex5 together with Eci1 and purified the complex. Employing Cryo-electron microscopy (Cryo-EM), we reconstructed the Pex5-Eci1 complex structure, uncovering a binding interface independent of PTS1 recognition.

Our work offers insight into a decades-old mystery of Eci1 targeting to peroxisomes. The identification of an additional binding interface between Pex5 and its cargo is crucial for understanding how proteins are targeted to peroxisomes with or without a PTS1. This highlights the intrinsic complexity of targeting to this multi-faceted and highly regulated metabolic organelle.

## Results

### Eci1 is targeted to peroxisomes when its PTS1 is masked by a fluorophore

In recent years, it is becoming apparent that targeting proteins to the peroxisome matrix by Pex5 is more complex than previously thought (Yifrach *et al*., 2016; Rymer *et al*., 2018). One example of this complexity is Eci1, which was shown to localize to peroxisomes even when its canonical PTS1 targeting signal was absent (Karpichev and Small, 2000; Yang, Purdue, and Lazarow, 2001). This PTS1 independent targeting led to the suggestion that Eci1 has a PTS2 and can use Pex7 in addition to Pex5, or that it is piggybacking on its paralog Dci1 (Geisbrecht *et al*., 1999; Karpichev and Small, 2000; Yang, Purdue and Lazarow, 2001). Nevertheless, how Eci1 is targeted to peroxisomes in the absence of a PTS1 had remained a mystery.

Intrigued by this unresolved question, we first masked the PTS1 of Eci1 by genomically integrating an mNG-encoding gene downstream *ECI1*. The synthesized protein is C’ tagged with mNG, preventing the ability of Pex5 to bind to the PTS1, which must be at the most C-terminus of a protein for functional recognition (Gould *et al*., 1989). In line with previous observations, we found that Eci1-mNG localizes to peroxisomes (Figure 1A). This is in stark difference to other PTS1-dependent peroxisomal proteins that lost their peroxisomal localization upon masking their PTS1 with mNG (Supplementary Figure S1) and more similar to Pox1, a well-studied Pex5 cargo that uses PTS1 independent targeting (Klein *et al*., 2002; Kempiński *et al*., 2020).

**Figure 1.**
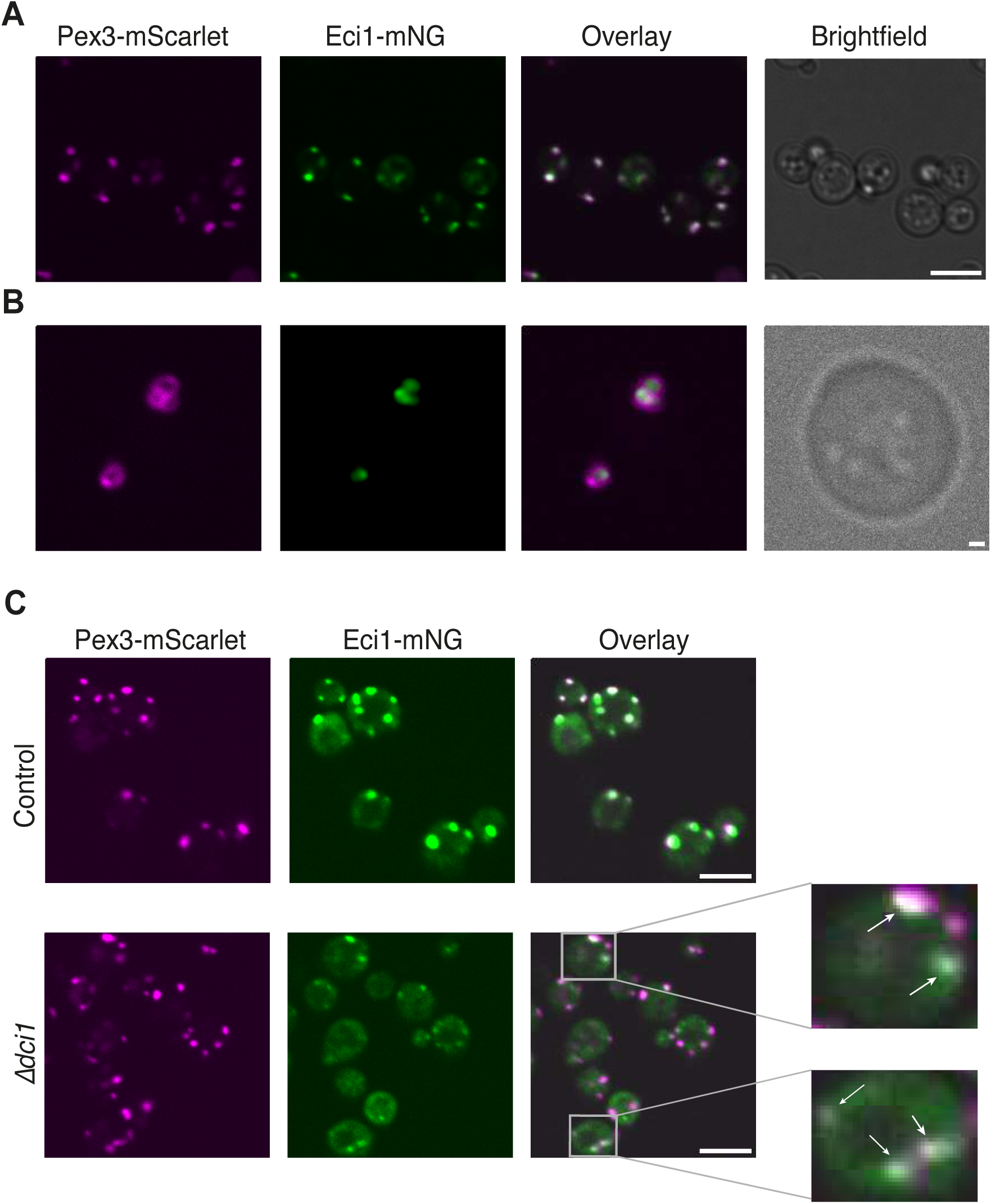
Eci1 is localized to peroxisomes when its PTS1 is masked by mNeonGreen, and its paralog is absent. A) Fluorescent microscopy images of Eci1-mNeonGreen (mNG) show co-localization with peroxisomes (Pex3-mScarlet) when C’ tagged. Scale: 5 μm. B) High-resolution (SORA) fluorescence microscopy images examining the sub-peroxisomal localization of Eci1-mNG on the background of *Δpex11* and following growth in oleate as a sole carbon source; Eci1 is in the peroxisomal matrix. Scale: 500 nm. C) Fluorescent microscopy images demonstrating that in the background of *Δdci1*, the signal of Eci1-mNG is heavily reduced, yet peroxisomal localization of Eci1 is still observed (indicated by white arrows). Scale: 5 μm.

The size of yeast peroxisomes is ∼150nm, which is lower than the diffraction limit of light (∼250nm). Hence, hypothetically, our microscopy images showing colocalization of Eci1-mNG and peroxisomes could be due to mislocalization to the surface of peroxisomes or proximal organelles. Hence, to verify that Eci1-mNG is indeed targeted to the peroxisome matrix, we increased the size of peroxisomes (by deletion of the *PEX11* gene and growth on oleate as a sole carbon source (Erdmann and Blobel, 1995; Yifrach *et al*., 2022) alongside imaging with a high-resolution microscopy system (SORA)). Indeed, we were able to affirm that Eci1-mNG is correctly localized to the peroxisomal matrix (Figure 1B). These results demonstrate that masking the PTS1 of Eci1 does not prevent its accurate targeting to the peroxisome matrix.

As previously suggested (Yang, Purdue and Lazarow, 2001), we considered that Eci1 could be “piggybacking” on its paralog, Dci1. Hence, we deleted *DCI1* and examined the effect on the peroxisomal localization on Eci1-mNG. Deleting *DCI1* resulted in impairment, but not complete abolishment, of the peroxisomal targeting of Eci1-mNG (Figure 1C). This implies the existence of an additional mechanism through which Eci1 can be targeted to peroxisomes, even when its PTS1 is masked, and its paralog is absent.

### Pex5 is the only cargo factor targeting Eci1

One possibility by which Eci1-mNG can be targeted to peroxisomes when Dci1 is absent, is by utilizing a cargo factor other than Pex5. To explore whether an alternate cargo factor can target Eci1 to peroxisomes when its PTS1 is manipulated, and Dci1 is absent, we tested several hypotheses.

Initially, it was suggested that Eci1 contains a PTS2 (Karpichev and Small, 2000). Hence, we deleted *PEX7*, the cargo factor responsible for targeting PTS2 proteins, either alone or on the genetic background of *Δpex5.* We observed that the targeting of Eci1 was unaffected when Pex7 was absent (Figure 2), contradicting the idea that Eci1 utilizes the Pex7-PTS2 targeting mechanism. Although Pex7 is not necessary for targeting, we wondered if it might still be sufficient for this process. To explore this possibility, we over-expressed Pex7 in the absence of Pex5 and could not see an effect. This shows that under the conditions that we performed our assay, Pex7 is neither necessary nor sufficient for Eci1-mNG targeting (Figure 2).

**Figure 2.**
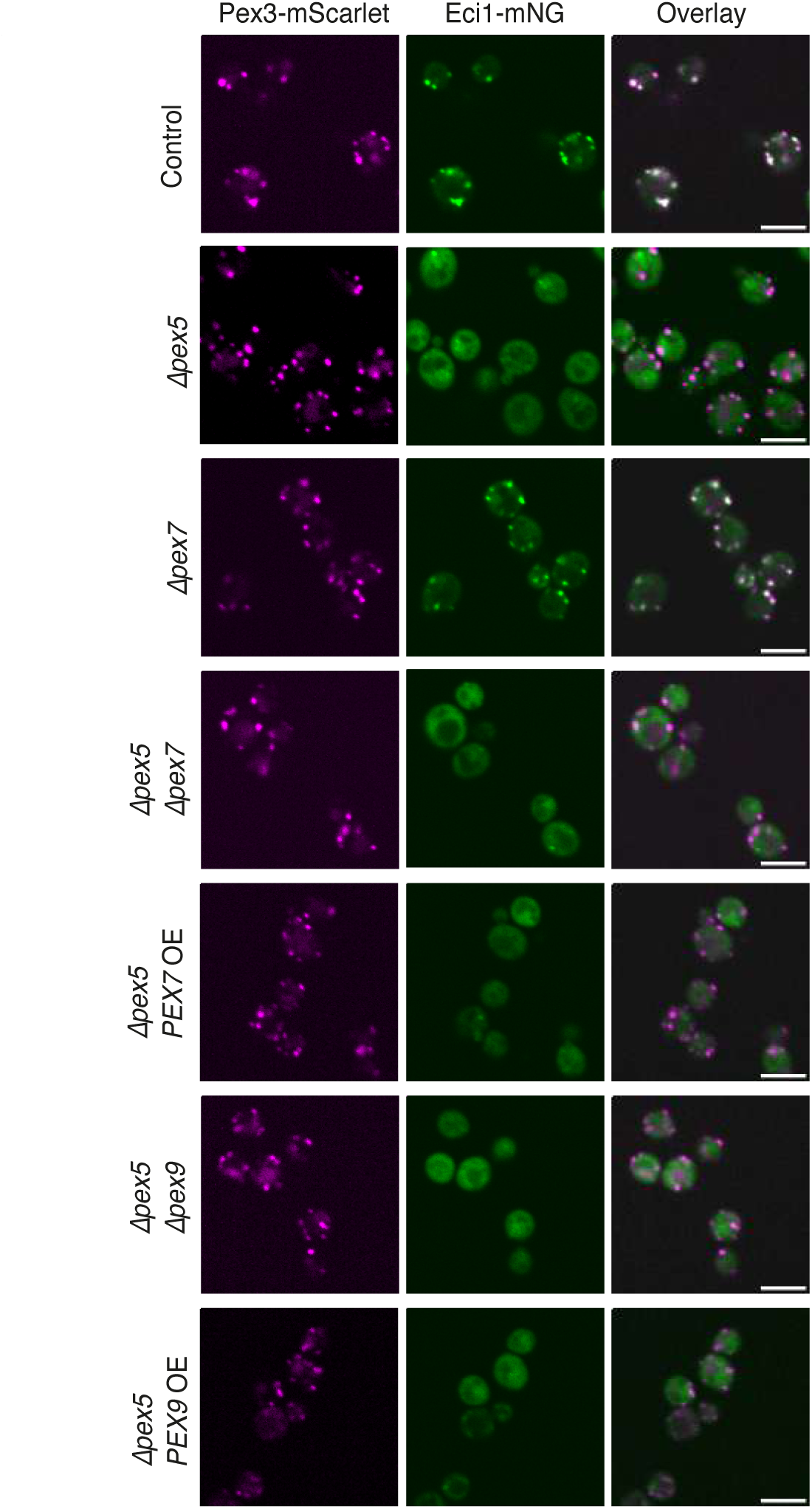
Eci1 is targeted solely by Pex5. Fluorescence microscopy images of different genetic backgrounds were examined to determine how Eci1 is targeted to peroxisomes and demonstrate a Pex5-dependent, Pex7 and Pex9-independent targeting mechanism. Scale: 5 μm.

Next, we examined the possibility that Eci1 can be targeted to peroxisomes by Pex9, which targets a specific set of PTS1 proteins to the peroxisomal matrix (Effelsberg *et al*., 2016; Yifrach *et al*., 2016, 2023). To examine the potential involvement of Pex9 in Eci1 targeting, we deleted *PEX9* (alone or on the background of *Δpex5*) or overexpressed it on the background of *Δpex5*. However, we did not observe any significant effect in either case. This again shows that Pex9 is neither sufficient nor necessary for targeting of Eci1-mNG. Taken together, we conclude that Pex5 is the only cargo factor that mediates Eci1 targeting to peroxisomes under the conditions that we were working.

### Eci1 directly binds Pex5 in the absence of its PTS1

To examine whether Eci1 can directly bind Pex5 in a PTS1-independent manner in the absence of Dci1 or any additional yeast protein, we employed an *in-vitro* co-expression system in *Escherichia coli (E. coli*). We expressed the Pex5 protein alongside Eci1 with (WT) or without (ΔPTS1) its most C terminal three amino acids (Figure 3). While deleting the PTS1 reduced the binding between Eci1 and Pex5, a direct binding was still observed. This suggests an additional interaction surface, that enables Pex5 to bind Eci1 in a PTS1-independent manner.

**Figure 3.**
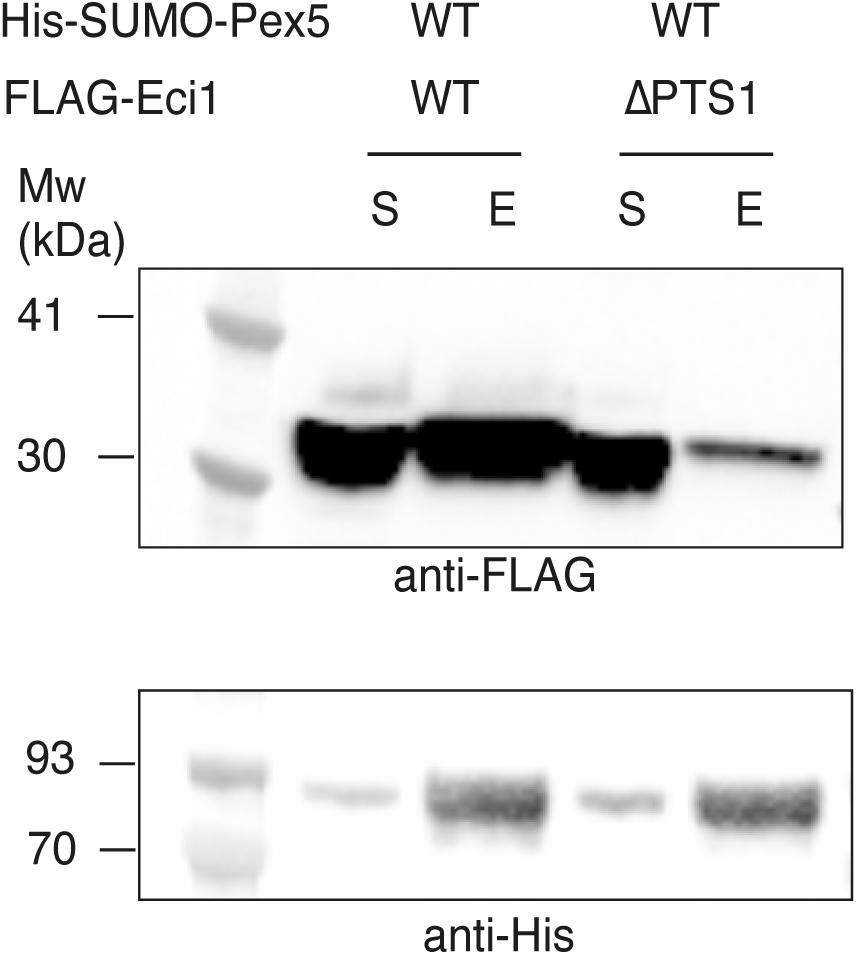
Eci1 can bind Pex5 in the absence of its PTS1. Western blot analysis of *in vitro* His-tag pull-down of Pex5 WT with either Eci1 or Eci1 without its Peroxisomal Targeting Sequence 1 (ΔPTS1). Blots were incubated with either anti-FLAG (upper blot) for the detection of FLAG-Eci1 or anti-His for the detection of His-SUMO-Pex5 (lower blot). When abolishing the PTS1, Eci1 exhibits a lower, yet clear, ability to bind Pex5 *in vitro*. S-soluble fraction, E-elution fraction.

### Eci1 binds to Pex5 through a non-canonical interface

To obtain a structure of the Pex5-Eci1 complex, we co-expressed full-length yeast Pex5 and Eci1 in *E. Coli*, purified and imaged them by single particle cryo-EM (Supplementary Figures S2-S5). Cryo-EM reconstruction of the Pex5-Eci1 complex dataset revealed Eci1 in a hexameric form, with a single subunit bound to a Pex5 monomer. The Eci1 hexamer was readily resolved to high resolution, whereas the bound Pex5 density was poorly resolved and appeared conformationally heterogeneous with respect to the Eci1. We, therefore, carried out two 3D variability analyses with the aim of determining the Eci1-Pex5 stoichiometry and conformational landscape (Supplementary Figures S2, S3, summarized in Supplementary Figure S4). First, variability analysis using a wide spherical mask around the complex indicated that the dataset primarily comprises singly bound Pex5 complexes, while a few subsets can be seen with two or more Pex5 densities bound to Eci1 subunits. Notably, all Pex5 densities adopt a similar orientation relative to the Eci1 hexamer (Supplementary Figure S2). Second, variability analysis using a mask to include the Eci1 hexamer and a single Pex5 indicated that the Pex5 binds with multiple conformations (Supplementary Figure S3). Refined 3D reconstructions from the separated subsets show that the Pex5 forms a variable interface with an Eci1 subunit. In some subsets, the interface includes only two attachment points with a gap in between, while in others, a more extensive, continuous interface is formed. The number of orientations that Pex5 adopts relative to Eci1 are restricted accordingly, so that Pex5 is poorly resolved when bound at two points only, and better resolved when the extensive interface is formed. Intermediate conformations are also seen, where partial density is formed in the gap.

To gain molecular insight, we focused on the subset in which Pex5 and the Eci1-Pex5 interface were most stable. The maps from this subset refined to 2.7 Å globally, while Pex5 and the interface were resolved to 2.9 Å following local refinement of this region (Supplementary Figures S4 and S5).

Looking at the atomic coordinates, the Eci1 is composed of an N-terminal core domain (M1-N200) with a spiral fold topology and a C-terminal region (M201-L280) that forms an α–helical trimerization domain. In the trimerization domain, three Eci1 subunits assemble into a characteristic trimeric disk with tight interactions between the three subunits. Two of these trimeric disks combine to form a hexamer (Figure 4A). Since the 3D refinement was done without applying symmetry, the six Eci1 subunits feature differences in the resolved residues. As such, the subunits contain residues I5-Q270, E4-Q270, I5-Q270, E4-Q270, and I5-Q270 (blue, purple, orange, cyan, and green in Figure 4A, respectively). The Pex5-bound subunit (in red, Figure 4A) contains a unique segment at its C-terminal end (^271^LGSKQRKHRL^280^) that is not observed in the other five subunits. Furthermore, this unique C-terminal segment has not been observed previously in any known apo-Eci1 structure. Within this segment, residues ^278^HRL^280^ delineate the PTS1 peroxisomal targeting signal, and this segment adopts a specific conformation influenced by its interaction with Pex5.

**Figure 4.**
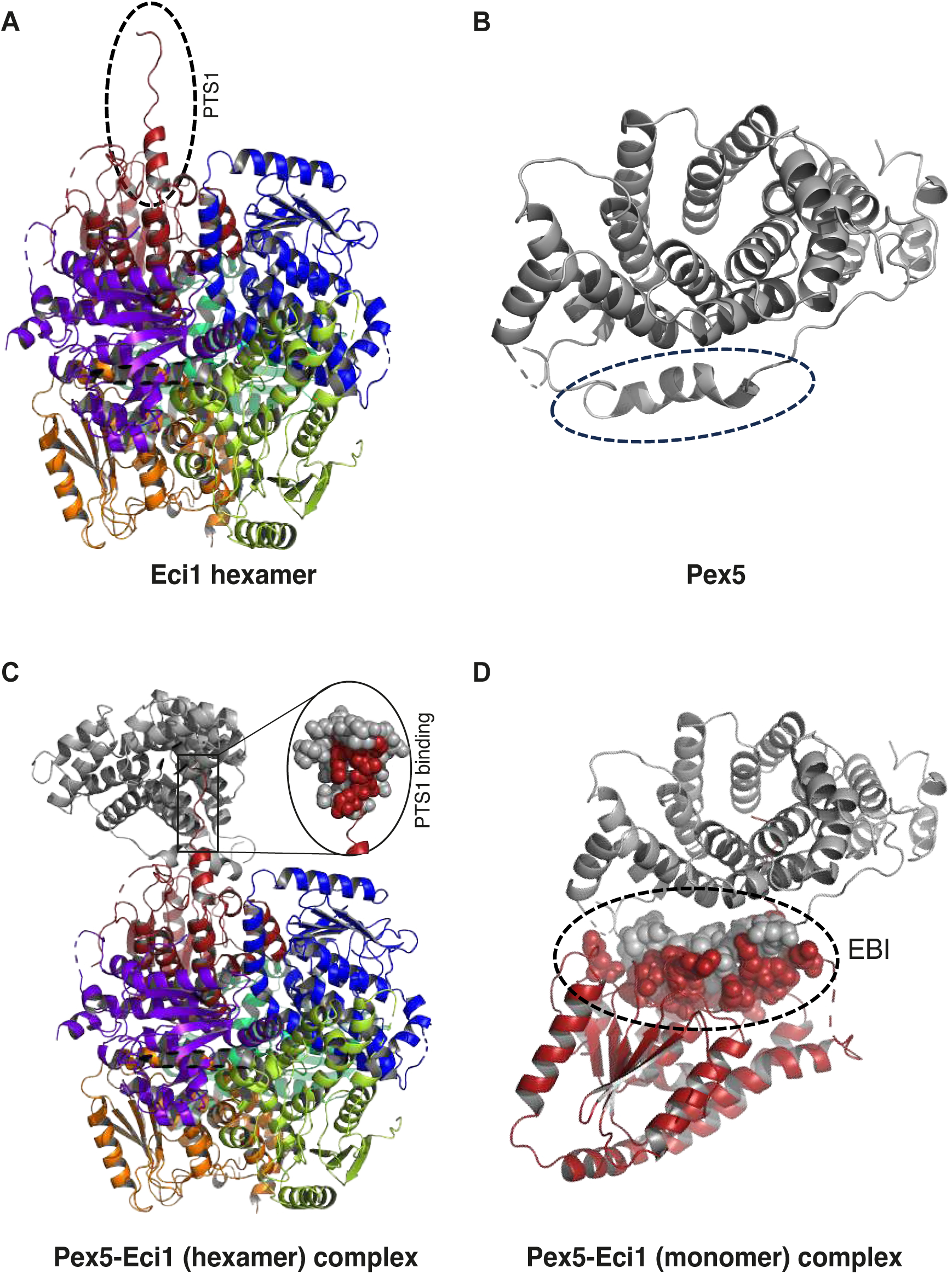
The Cryo-EM structure of Pex5 in complex with an Eci1 hexamer highlights a PTS1 binding interface and a novel Eci1 binding interface (EBI). A) The Eci1 hexamer consists of six monomers shown in ribbon and represented in various colors. A black dashed circle represents the additional 10 residues at the C-terminal region (^271^LGSKQRKHRL^280^) of the red monomer that have not been observed previously. B) The Pex5 cargo factor comprises the TPR domain represented in a grey ribbon. The additional 21 residues at the N-terminal region (^276^LVNDDLNLGEDYLKYLGGRVN^296^) have not been observed previously (blue dashed circle). C) The C-terminal domain of Pex5 is represented as a grey cartoon. The Eci1 hexamer interacting with Pex5 is shown in red; the binding interface involving the C-terminal segment of Eci1, including the PTS1 (^278^HRL^280^), represented as red balls, is interacting with the TPR3 domain of Pex5. D) The novel EBI of Pex5 with Eci1 is an elongated stretch of 21 mostly conserved residues in the N-terminal domain (NTD) of Pex5, represented as grey balls interacting with highly conserved residues on the surface of Eci1 as indicated by red balls; the newly identified binding interface is highlighted in a black dashed circle.

The Pex5 structure (L276-F612) comprises the TetratricoPeptide Repeat (TPR) domain (N314-S553) with seven TPR repeats, each with 34 residues (α1−α2, α3−α4, α5−α6, α7−α8, α9−α10, α11−α12, and α13−α14) (Supplementary Figure S6 and represented as a cartoon in Figure 4B). This structure also includes the PTS1-cargo binding region, which is a known feature of the Pex5 receptor TPR domain. Notably, there is electron density for an additional 21 residues at the N-terminal region (^276^LVNDDLNLGEDYLKYLGGRVN^296^) that have not been observed previously. The C-terminal region of the red Eci1 monomer interacts with specific residues from the TPR3, TPR6, and TPR7 repeats of Pex5. The Cryo-EM maps of the Pex5-Eci1 complex containing the full-length Pex5 do not show electron density for the N-terminal domain (NTD) of Pex5 (up to E275). This indicates that the NTD of Pex5 is flexible in the context of the complex, making it challenging to visualize using Cryo-EM.

Importantly, the Cryo-EM structure of the Pex5-Eci1 complex (depicted in Figures 4C and 4D) reveals an extensive protein-protein interface facilitated by two binding surfaces. The first involves the C-terminal segment (^272^GSKQRKHRL^280^) of one Eci1 subunit, incorporating the PTS1 signal (^278^HRL^280^), which interacts with the TPR3 and TPR6 domains of Pex5 (Figure 4C). The segment encompassing the PTS1 residues is situated within a deep central cavity of the TPR domain. This interaction is reinforced by a network of hydrogen bonds formed between ^278^HRL^280^ and residues in TPR3, TPR6 and TPR7 of Pex5. Among these interactions is a salt bridge between R279 of Eci1 and E394 of Pex5. Additional interactions involve the C-terminal segment of Eci1, specifically ^272^GSKQRKHRL^280^, with residues in the TPR3 domain of Pex5 and a few residues (G292, V295 and N296) from the additional N-terminal region of Pex5 (Figure 4C).

Importantly, our structural analysis revealed an additional, newly discovered, Eci1 binding interface (EBI) involving an elongated stretch of 21, mostly conserved, residues in the NTD of Pex5 that have not been previously observed (^276^LVNDDLNLGEDYLKYLGGRVN^296^, Figure 4B, Supplementary Figure S6). This stretch is involved in binding a distinct interface in Eci1. This observation suggests that the EBI and the PTS1 binding interface operate independently and complementarily in mediating the interaction between Pex5 and Eci1.

Focusing on the newly identified EBI, we observed that it interacts with highly conserved residues on the surface of Eci1 (Figure 4D). Within this binding interface, two segments of Eci1, ^65^FFSSGADFKGIAK^77^ and ^143^KVYLLYP^149^, play important roles in forming core interactions with the NTD of Pex5. These interactions involve the formation of hydrogen bonds and salt bridges, thereby enhancing the stability and specificity of the Pex5-Eci1 interaction. To estimate the evolutionary conservation of amino acids within Eci1 and related proteins, we used the ConSurf server (Ashkenazy *et al*., 2016), which generates multiple sequence alignments of 150 homologous proteins and predicts the conservation of amino acids based on their evolutionary history. The algorithm clearly detects the high conservation among the amino acids interacting with Pex5 (represented in maroon in Supplementary Figure S7), supporting the importance of these residues in facilitating the Pex5-Eci1 interaction.

Both binding interfaces exhibit shape complementarity, with the concave surface of Eci1 fitting well with the convex surface of Pex5 (Figure 5A). In addition, there is electrostatic complementarity between the mostly electronegative surface (highlighted in red) of Pex5 and the mostly electropositive surface (highlighted in blue) of Eci1 (Figure 5B).

**Figure 5.**
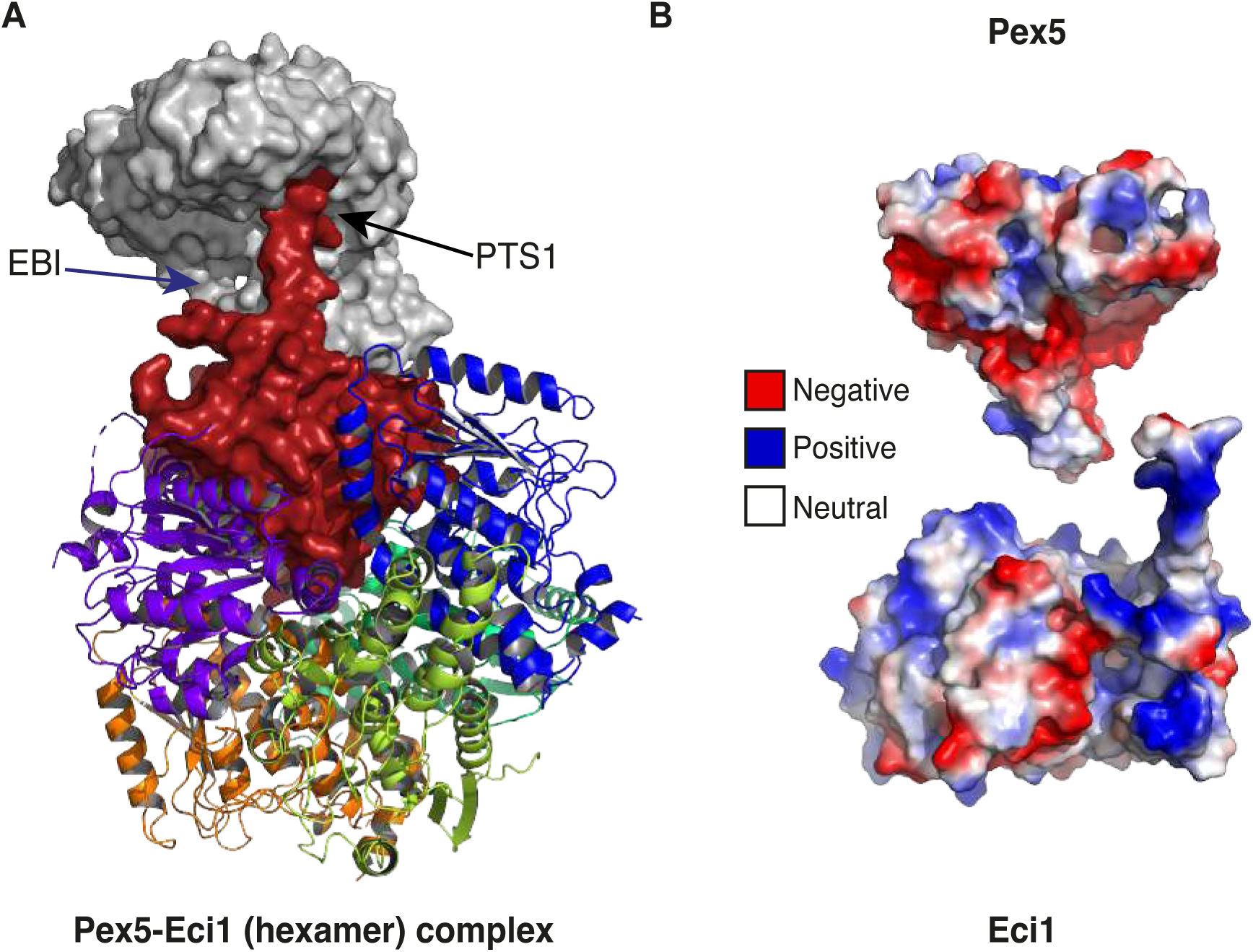
The Pex5-Eci1 complex structure interfaces exhibit shape and charge compatibility. A) Surface representation of Pex5 (grey) interacting with one subunit of the Eci1 hexamer (red) and ribbon representation of the other five Eci1 monomers. Two binding interfaces were identified, one involving the C-terminal PTS1 of Eci1 interacting with the TPR domain of Pex5 (black arrow) and the newly identified EBI (blue arrow). Both interfaces exhibit shape complementarity. B) Electrostatic representation of the electropositive surface of Eci1 monomer (bottom), which is complementary with the electronegative surface of Pex5 (top). Electronegative surfaces are depicted in red, electropositive in blue, and neutral in white.

The formation of the complex also relies on multiple salt-bridge interactions between specific amino acid residues in both proteins that play a critical role in stabilizing the complex. Specifically, negatively charged residues in Pex5; N298, N393, E394, E361 and D397 engage in salt bridges with positively charged residues in Eci1; K73, R279, and R276. Additionally, a salt bridge forms between the negatively charged residues N29 and D71 in Eci1 and the positively charged residue R294 in Pex5. Notably, R276 of Eci1 participates in multiple salt bridge interactions with D397 of Pex5, and R279 of Eci1 is involved in salt bridge contacts with E361, N393, and E394 of Pex5. This interplay of oppositely charged residues contributes to the overall stability and specificity of the Pex5-Eci1 interaction.

The synergy of both shape complementarity and electrostatic complementarity, exemplified by salt bridge interactions, amplifies the binding strength between Pex5 and Eci1. These characteristics guarantee a precise and secure interaction between the proteins, elucidating the precise targeting even in the absence of the PTS1 of Eci1.

### The N-terminal segment of Eci1 has a dual role

Eci1 catalyzes a critical step in fatty acid metabolism, facilitating the oxidation of unsaturated fatty acids. Specifically, it catalyzes the conversion of 3E- and 3Z-enoyl-CoA thioesters to 2E-enoyl-CoA thioesters, which are intermediates in the four-step oxidation pathway. For this role, it must bind Coenzyme A (CoA). The structure of Eci1 with CoA was previously solved (PDB entry 4ZDB) (Onwukwe *et al*., 2015). Structural alignment between the Eci1-Pex5 complex and the Eci1-CoA complex revealed that the newly defined NTD segment of Pex5 (^276^LVNDDLNLGEDYLKYLGGRVN^296^) occupies the same position typically occupied by CoA in Eci1-CoA complex (Supplementary Figure S8). The binding of Eci1 to both CoA and Pex5 is mediated by the same residues, namely D71, F72, and S68. This shared interaction raises the possibility of either competition or coordination between CoA and Pex5 for binding to Eci1.

### Pex5 interacts with different cargos via distinct binding interfaces

Our discovery of an additional binding interface on Pex5 for a protein that already contains a PTS1 is not unique. In fact, this has previously been shown for human PEX5 binding to the human Alanine-glyoxylate aminotransferase (AGT) (Fodor *et al*., 2012). AGT is a peroxisomal enzyme that contains PTS1 and exhibits an extensive interface with PEX5 also via additional, non-PTS1 interactions. We, therefore, set out to compare whether Pex5 binding to Eci1 and PEX5 binding to AGT occur through similar interfaces.

We performed a structural alignment and interaction comparison between the TPR domain of yeast Pex5 with Eci1 and human PEX5 with AGT (PDB entry 3R9A (Fodor *et al*., 2012)). Despite a relatively low sequence identity (35%), the TPR domain of Pex5 in yeast and humans exhibited a high degree of structural similarity with a root mean square deviation (RMSD) of 0.858 Å (Figure 6A). However, AGT forms 29 hydrogen bond contacts with PEX5, whereas Eci1 forms 97 contacts with Pex5. We first focused on the PTS1 region, which aligns perfectly (Figure 6B and 6C). We observed that the PTS1 of both proteins interacts with conserved residues in Pex5/PEX5 (N393, A502, N503, N530 in Pex5 and N415, A533, N534 and N568 in PEX5 respectively), indicating the fundamental role of PTS1 in PEX5 interactions.

**Figure 6.**
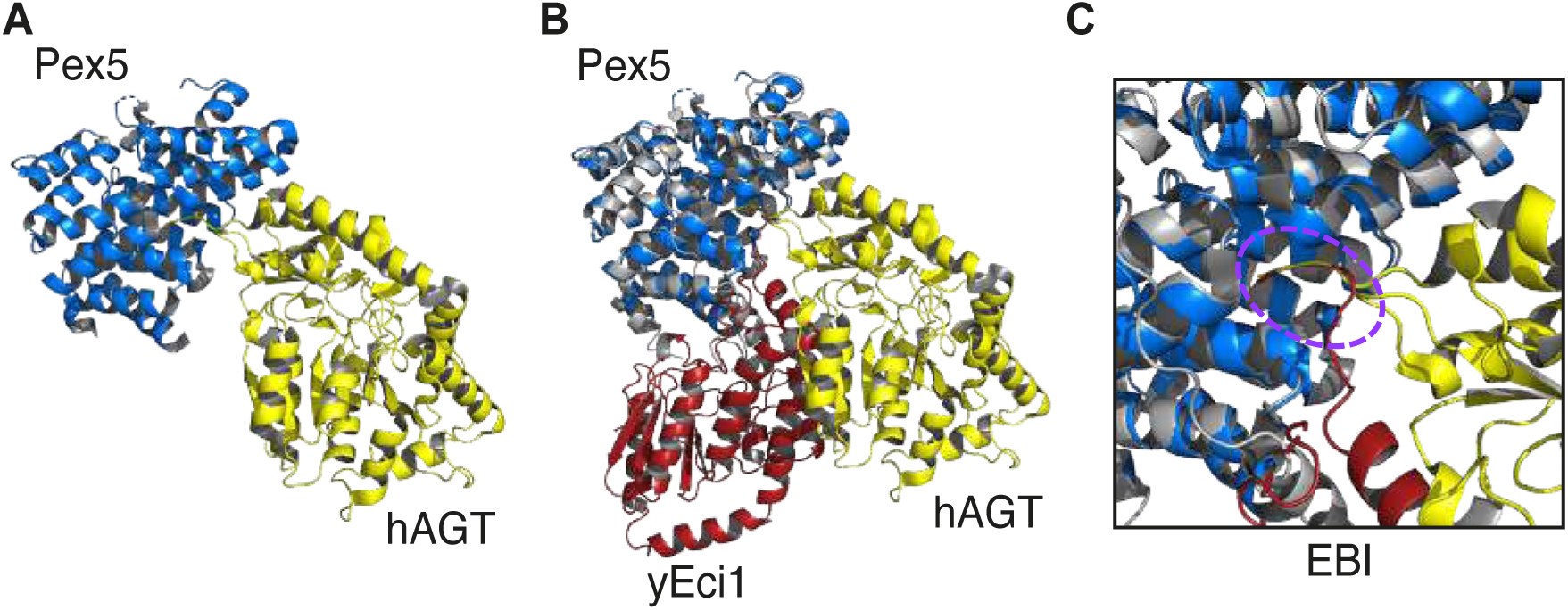
The newly defined Pex5-Eci1 EBI differs from the binding interface of PEX5 with AGT. A) The human PEX5-AGT complex is depicted in blue and yellow, respectively (PDB entry 3R9A). B) The yeast Pex5-Eci1 complex depicted by grey and red ribbons respectively, aligned with the PEX5-AGT structure. Notably, the TPR domain of yeast Pex5 and human PEX5 exhibits a strong alignment. There is a resemblance observed in the PTS1 binding interface between Eci1 and AGT (depicted by the dashed purple circle in panel C) The second binding interface is located in different regions within the Pex5/PEX5 structure and involves distinct residues.

We further analyzed the additional binding interface of each pair. In the PEX5-AGT interaction, the second binding interface is localized and relatively weak, involving only three residues, while in the Pex5-Eci1 interaction, the second interface is more extensive, engaging 12 residues in Pex5 and 15 residues in Eci1 (Figure 6). Importantly, the amino acids on the Pex5/PEX5 side are distinct with Pex5 interacting in N278, D279, D280, L281, L283, G284, Y287, Y290, R294, V295, N296 and N298 PEX5 in R608, L609 and S612. This highlights that Pex5 possesses unique binding interfaces tailored to different binding partners, potentially offering adaptability to enable cargo-specific targeting.

The existence of separate binding interfaces for various cargo implies the potential existence of additional cargo proteins that may be targeted in a PTS1 independent manner. To investigate this possibility, we employed AlphaFold2 to model the structure of Dci1 and aligned it with our established structure of the Pex5-Eci1 complex. Remarkably, many residues of Eci1 that interact with Pex5 are highly conserved in its paralog Dci1 (Supplementary Figure S9). Specifically, residues F65, S68, F72, P122, Y145, P149, P183, F268 and L271 in Eci1 correspond to Y59, S62, F66, P116, F139, P143, P147, P177, F259 and L262 in Dci1, respectively. This suggests the intriguing possibility that Dci1 might utilize a similar, non-canonical peroxisomal targeting pathway via Pex5.

## Discussion

While the role of the PTS1 of Eci1 in binding to Pex5 and mediating targeting is unrefuted, it has been a longstanding mystery as to how Eci1 can be targeted to peroxisomes in the absence of its PTS1 (Geisbrecht *et al*., 1998, 1999; Gurvitz *et al*., 1998; Karpichev and Small, 2000; Yang, Purdue and Lazarow, 2001). In this work, we show that the targeting of Eci1 to the peroxisomal matrix is exclusively dependent on Pex5 and is not mediated by Pex7 nor by Pex9. We also showed that Eci1 targeting in the absence of its PTS1 cannot be explained solely by piggybacking on its paralog Dci1. Our *in vivo* findings implied that Eci1 is targeted to peroxisomes in a Pex5-dependent, PTS1-independent manner, even when Dci1 is absent. These findings suggested an additional Pex5 binding site in Eci1, leading us to solve the structure of Pex5-Eci1 using Cryo-EM.

The structures of the TPR domains of human Pex5 with either AGT or SCP2 were previously solved (Williams *et al*., 2011; Fodor *et al*., 2012). Recently, the structure of full length *Trypasonoma cruzi* Pex5 and Malate dehydrogenase (MDH) was also solved (Sonani *et al*., 2023). Here, we co-expressed the full-length yeast Pex5 and Eci1, allowing us to resolve and visualize never-before-seen residues in both proteins and uncover a novel binding interface between them, in line with our *in vivo* findings.

Why has this newly identified EBI with Eci1 not been discovered so far? Previous studies in *Saccharomyces cerevisiae* mutated residue N495 in Pex5 to argue that all Eci1 binding is through the PTS1 binding site. This point mutation was previously considered to abolish only the PTS1 binding domain. However, structural data of Pex5 (including in this study) suggest that mutating this crucial residue within the core of Pex5 can not only abolish PTS1 but destabilize the entire protein structure, which can possibly lead to a non-functional Pex5 and misinterpretation of the effects of this point mutation.

The findings in this study offer the intriguing possibility that other peroxisomal matrix proteins are targeted by Pex5 in a PTS1-independent manner (for example, Supplementary Figure S9). In recent years, many proteins have been suggested to interact and bind to Pex5 in a non-canonical manner, such as the Oxalyl-CoA synthetase Pcs60 in yeast, which has been shown to still bind Pex5 even when it was lacking its PTS1 (Bürgi *et al*., 2023). Another example is the Acyl CoA oxidase Pox1, which has been suggested to harbor a noncanonical PTS as a signal patch in the fully folded protein (Klein *et al*., 2002; Kempiński *et al*., 2020). This phenomenon extends to other organisms, including observations of the human AGT protein (Fodor *et al*., 2012), which also interacts with Pex5 in a PTS1 independent manner. A similar observation was shown in *Arabidopsis thaliana,* with Catalase (CAT2) being targeted to peroxisomes in the absence of a PTS1 (Al-Hajaya *et al*., 2022). Recently, it was shown that in *Trypasonoma cruzi*, Malate dehydrogenase (MDH) has a noncanonical interaction with Pex5 when in complex with Pex14 (Sonani *et al*., 2023), where the N terminal part of Pex5 is an intrinsically disordered region (IDR) as in this study (data not shown). Furthermore, this highlights the importance of using a full-length Pex5 for targeting studies, as different parts of its N-terminal domain are likely to become structured upon binding different proteins. This binding stabilizes the IDR and allows for a more detailed resolution of the structure. Interestingly, in two of the cargos mentioned above-Pcs60 hexamer and MDH tetramer-Pex5 only partially occupies the available binding interfaces on the cargo, (Bürgi *et al*., 2023; Sonani *et al*., 2023). This is in line with our data, where only one Eci1 monomer of the hexamer binds subunit is observed bound to Pex5 in most complexes (Supplementary Figure S2). Further study is required to determine the nature of this observation, focusing primarily on whether this phenomenon also occurs *in vivo* and its possible functional importance for peroxisomal targeting.

Moreover, systematic analysis in yeast has uncovered many Pex5 cargo proteins that are Pex5-dependent but PTS1-independent (Yifrach *et al*., 2022). These numerous findings beg a paradigm shift for Pex5 targeting as we have conceived it to date; further study is required to elucidate the biological importance of non-canonical Pex5 binding interfaces. Leaving aside the “canonical” mindset by which we have been studying peroxisomal targeting so far will allow us to find new peroxisomal proteins that use non-canonical interaction for their targeting. Moreover, it will enable to shed light on the dynamic manner by which Pex5 operates in targeting of many proteins to peroxisomes, allowing several levels of regulation using multiple binding sites, including prioritizing some cargos over others (Rosenthal *et al*., 2020) in line with the needs of the cell and organism.

## Materials and methods

### Yeast strains and strain construction

All strains in this study were based on the strain yMS4097. Genetic manipulations were performed using a homologous recombination-based transformation of a suitable PCR product using the lithium-acetate method (Wei Xiao, 2006). The correct insertion of the PCR product was verified in all strains. The primers in this study used for yeast genetic manipulation were designed using the web tool Primers-4-Yeast (Yofe and Schuldiner, 2014). All strains, primers, and plasmids in this study are listed in the supplementary tables 1,2,3 respectively.

### Yeast growth media

The synthetic media used in this study contains 6.7 g/L yeast nitrogen base with ammonium sulfate (Conda Pronadisa #1545) and 2% glucose (SD) or 0.2% oleic acid (Sigma-Aldrich, St. Louis, MO, USA) + 0.1% Tween 80 (Sigma-Aldrich, St. Louis, MO, USA) (S-oleate), both with a complete amino acid mix (oMM composition) (Hanscho *et al*., 2012). When Geneticin antibiotic was used, the media contained 0.17 g/L yeast nitrogen base without ammonium sulfate (#1553 Conda Pronadisa, Madrid, Spain) and 1 g/L of monosodium glutamic acid (Sigma-Aldrich, St. Louis, MO, USA #G1626). The strains were selected using a dropout mix (same composition as “SD” above, without the specific amino acid for selection) or with antibiotics using the following concentrations: 500 mg/L Geneticin (G418; Formedium), or 200 mg/L Nourseothricin (NAT, WERNER BioAgents “ClonNat”).

### Microscopy

#### Manual microscopy

Images in Figures 1A, 1C, and 2 were obtained using manual microscopy as follows: Yeast strains were grown overnight in an SD-based medium with appropriate selection in 96-well polystyrene plates and were then transferred to S-oleate for 20 hours. The cultures in the plates were then transferred manually into glass-bottom 384-well microscope plates (Matrical Bioscience) coated with Concanavalin A (Sigma-Aldrich). After 20 minutes, the wells were washed twice with double-distilled water (DDW) to remove non-adherent cells and obtain a cell monolayer. Imaging was performed in DDW. The images were acquired using the VisiScope Confocal Cell Explorer system, composed of a Zeiss Yokogawa spinning disk scanning unit (CSU-W1) coupled with an inverted Olympus microscope (IX83; ×60 oil objective; Excitation wavelengths of 488 nm for mNeonGreen and 561 nm for mScarlet). Images were taken by a connected PCO-Edge sCMOS camera controlled by VisiView software. for all micrographs, a single, representative focal plane was shown.

### High-resolution imaging (SORA)

A strain containing a *PEX11* deletion (Figure 1B) was grown in S-oleate and prepared for imaging in the depicted manner above (“Manual microscopy”). Images were taken using the Olympus IXplore SpinSR system, composed of an Olympus IX83 inverted microscope scanning unit (SCU-W1) in addition to a high-resolution spinning disk module (Yokogawa CSU-W1 SORA confocal scanner with double microlenses and 50 µm pinholes), operated by ScanR. Cells were imaged using an X60 oil lens (NA 1.42) and Hamamatsu ORCA-Flash 4.0 camera. Images were recorded in two channels: mNeonGreen (excitation wavelength 488 nm) and mScarlet (excitation wavelength 561 nm). Cells were imaged using Z-stacks (19 stacks, 0.2μm distance between the stacks); for all micrographs, a single, representative focal plane was shown.

### Eci1 and Pex5 plasmid construction

All cloning reactions were performed by the Restriction-Free (RF) method (Unger *et al*., 2010) Full-length *PEX5* (1-612) from *Saccharomyces cerevisiae* was cloned into the expression vector pET28-bdSumo (Zahradník *et al*., 2019)), using Restriction Free cloning (Unger *et al*., 2010) yielding Pex5 with a cleavable N-terminal His-Sumo fusion. Yeast *ECI1* was cloned into the first open-reading frame of the expression vector pACYCDuet-1 (Novagen), which includes an N-terminal Flag-tag followed by a TEV cleavage site. The ΔPTS1 Eci1 construct was generated by the deletion of the last three amino acids (^278^HRL^280^) of the protein. The reaction was performed by Transfer-PCR (TPCR) (Erijman *et al*., 2011). Primers used for cloning of WT Eci1, WT PEX5 and for generation of ΔPTS1 Eci1 are listed in Table S2.

### Eci1 and Pex5 protein co-expression

*ECI1* (in *pACYCDut-FLAG-Eci1*) and *PEX5* (in *pET28-bdSumo-PEX5*) were co-expressed in *E. coli* BL21(DE3). Expression was performed in LB medium supplemented with the appropriate antibiotics (Kanamycin and chloramphenicol). For small-scale and large-scale expression, 10ml and 5L cultures, were used, respectively. Expression was induced with 200μM IPTG followed by shaking at 15 °C for ∼18 hours. Cell pellets were stored at -20 °C before processing.

### Small-scale Eci1 and Pex5 protein pull-down

Cell pellets (derived from 10 ml culture) were lysed by sonication in Tris-buffered saline (TBS) buffer supplemented with 1 mM phenylmethylsulfonyl fluoride (PMSF) and 1μl/mL of protease inhibitor cocktail (Set IV, EMD Chemicals, Inc). Protein pull-down experiments were performed using Ni-resin (Adar Biotech) according to the manufacturers’ recommendations. Western blot analysis was performed using THE^TM^ DYKDDDDK Tag Antibody (HRP-conjugated; A01428, GenScript) and Monoclonal Anti-polyHistidine−Peroxidase (A7058, Sigma). Proteins were analyzed on 4-20% SurePAGE precast gels (M00657, GeneScript).

### Purification of Pex5-Eci1 complex

A cell pellet from a 5L culture was resuspended in lysis buffer (50mM Tris pH=8, 0.5M NaCl) supplemented with 200KU/100 ml lysozyme, 20ug/ml DNase, 1mM MgCl_2_, and protease inhibitor cocktail. The resuspended cells were lysed by a cooled cell disrupter (Constant Systems). The clarified lysate was incubated with 5ml washed Ni beads (Adar Biotech) for 1h at 4°C, after which it was loaded on GE column 16/20 connected to Aukta Purifier. After removing the unbound supernatant, the beads were washed with lysis buffer, followed by a stepwise washing with lysis buffer of 20 mM, then by a wash with 50 mM of imidazole. The beads were then equilibrated with sumo-cleavage buffer (40mM Tris pH=7.5, 250mM NaCl, 250mM Sucrose, 2mM MgCl2). The Pex5-Eci1 complex was eluted from the washed beads by incubation of the beads with 5ml sumo-cleavage buffer (supplemented with 0.2mg *bdSumo* protease) overnight at 4°C. The supernatant fraction, containing the cleaved Pex5-Eci1 complex was removed, concentrated and applied to a size exclusion column (Superdex 200 10/300 GL, Cytiva) equilibrated with PBS. The Pex5-Eci1 complex, migrating as a single peak at 11ml, corresponded to a molecular weight >200kDa and <443kDa (based on the migration positions of amylose and apoFerritin on the same column, respectively). The presence of both Pex5 and Eci1 in the complex peak was verified by SDS PAGE stained with Coomassie brilliant blue. Eci1 coming along with Pex5 was also visualized by western blot analysis developed with THE^TM^ DYKDDDDK Tag Abs (data not shown). The pure complex was flash frozen in aliquots using liquid nitrogen and stored at -80°C.

### Sample preparation for EM

2.5 µl of Pex5-Eci1 complex at 2 mg/ml concentration was transferred to glow discharged Au-Flat 1.2/1.3 300 mesh grids (Protochips), blotted for 3 seconds at 4°C, 100% humidity, and plunge frozen in liquid ethane cooled by liquid nitrogen using a Vitrobot plunger (Thermo Fisher Scientific).

### Cryo-EM image acquisition

Cryo-EM data were collected on a Titan Krios G3i transmission electron microscope (Thermo Fisher Scientific) operated at 300 kV. Movies were recorded on a K3 direct detector (Gatan) installed behind a BioQuantum energy filter (Gatan), using a slit of 15 eV. Movies were recorded in counting mode at a nominal magnification of 105,000x, corresponding to a physical pixel size of 0.842 Å. The dose rate was set to 19.4 e-/pixel/sec, and the total exposure time was 1.6 sec, resulting in an accumulated dose of 45.5 e-/Å2. Each movie was split into 47 frames of 0.034 sec. Nominal defocus range was -1.0 to -1.5 μm, however the actual defocus range was larger. Imaging was done using an automated low dose procedure implemented in SerialEM v3.9-beta7 (Mastronarde, 2005). A single image was collected from the center of each hole using image shift to navigate within hole arrays containing up to 5x5 holes, and stage shift to move between arrays. Beam tilt was adjusted to achieve coma-free alignment when applying image shift.

### Cryo-EM image processing

Image processing was performed using CryoSPARC software v3.0.1 (Punjani *et al*., 2017). A total of 5,577 acquired movies were subjected to patch motion correction, followed by patch CTF estimation (Supplementary Figure S4). Of these, 4,727 micrographs having CTF fit resolution better than 5 Å and relative ice thickness lower than 1.2, were selected for further processing. Initial particle picking was done using the ‘Blob Picker’ job on a subset of micrographs. Extracted particles were classified in 2D and selected class averages showing the Eci1 hexamer features were used as templates for automated particle picking from all selected micrographs, resulting in 2,170,575 picked particles. Particles were extracted, binned 4x4 (64-pixel box size, 3.37 Å/pixel), and cleaned by multiple rounds of 2D classification, resulting in 1,083,831 particles. These particles were re-extracted, binned 2x2 (200-pixel box size, 1.68 Å/pixel) and used for *ab initio* 3D reconstruction with a single class, followed by non-uniform refinement. Both *ab initio* and non-uniform refinements revealed a single Pex5 monomer bound to the Eci1 hexamer. Using 3D variability analysis (Punjani and Fleet, 2021) (8 modes, 10 Å low-pass filter) with a spherical mask of 160 Å diameter imposed, followed by classification into 20 3D classes, a minor population of Eci1 hexamers bound to a varying number of Pex5 monomers was resolved (Supplementary Figure S2). In these subsets, bound Pex5 monomers could be refined to limited resolution due to conformational variability. Blurred density associated with multiply bound Pex5 monomers can also be observed in 2D class averages (Supplementary Figure S4). To better resolve Pex5 conformational variability and its interface with Eci1, we focused on the complexes of 6 Eci1 to 1 Pex5, which constituted the vast majority of the particles. 3D variability analysis was performed with imposed real-space solvent mask generated by the non-uniform refinement above (3 modes, 10 Å low-pass filter), followed by classification into 10 3D classes. Particles from all classes were re-extracted without binning into separate datasets, and subjected to non-uniform refinement. Subsequently, local refinement was performed using a soft-edged ellipsoid mask around Pex5 (including the Pex5-Eci1 interface) in initial iterations and a ‘dynamic’ solvent mask in final iterations, once resolution better than 5 Å was reached. This analysis clearly shows conformational variability in Pex5 binding (Supplementary Figure S3. Pex5 and the interface were best resolved in one of the classes (Supplementary Figure S4, 117,945 particles), which refined to 2.79 Å using non-uniform refinement and local refinement respectively. This class was used for downstream atomic coordinate modelling. The rest of the classes showed lower rigidity at the Pex5 area and Eci1 interface.

Particle images of smaller complexes were identified and processed, however their reconstructions were limited to low resolution, which made interpretation unreliable. Based on size and shape, these particles can be attributed to Pex5 or Eci1 monomers. 3D visualization was performed using UCSF Chimera (Pettersen *et al*., 2004a).

### Model building of Pex5-Eci1 complex

Model budling of the yeast Pex5-Eci1 complex included the docking of the known Eci1 hexamer crystal structure and the predicted model of full length Pex5 obtained using the AlphaFold2 software (Senior *et al*., 2020; Jumper *et al*., 2021) onto the Cryo-EM maps. The N-terminal segment of Pex5 (up to Y301) was predicted to be unstructured, with low accuracy predictions and confidence, as indicated by correspondingly low model prediction on the local distance different test (pLDDT) below 50. Due to this uncertainty, the N-terminal segment was truncated in the model. The predicted model of Pex5 (A302-F612) obtained from AlphaFold2 and the known structure of the Eci1 hexamer from *Saccharomyces cerevisiae* (PDB entry 1PJH) (Mursula, Hiltunen and Wierenga, 2004) were used as structural models for docking into the Cryo-EM maps, using the Dock-in-Map program in PHENIX (Adams *et al*., 2010). Specifically, the Eci1 hexamer was docked into the Pex5-Eci1 complex map and the Eci1-Pex5 complex into the Pex5 local map.

This integration allows for the reconciliation of computational predictions with experimental data, providing a comprehensive and accurate depiction of the Pex5-Eci1 complex structure. All steps of atomic refinements were carried out with the Real-space refinement in PHENIX (Klaholz, 2019). The Eci1 hexamer model was built into the Pex5-Eci1 complex map, and the Eci1-Pex5 model into the Pex5 local map using the COOT program (Emsley and Cowtan, 2004). The two models were evaluated with the MolProbity program (Chen *et al*., 2010).

The Cryo-EM map of the Eci1 hexamer shows electron density for I5-Q270, E4-Q270, I5-Q270, E4-Q270, and I5-Q270 in five copies, and L4-L280 for the Eci1 copy binding Pex5. This latter subunit contains a unique segment at its C-terminal end (^271^LGSKQRKHRL^280^) that is not observed in the other five subunits.

The Pex5 local refinement map of the Pex5-Eci1 complex containing the full-length Pex5 does not show electron density for the N-terminal domain (NTD) of Pex5 (up to E275) (Supplementary Figure S6). This indicates that the NTD of Pex5 is likely flexible or dynamic in the context of the complex, making it challenging to visualize using Cryo-EM. The Pex5 structure (L276-F612) comprises the TPR domain (N314-S553) with seven TPR repeats, each with 34 residues (α1−α2, α3−α4, α5−α6, α7−α8, α9−α10, α11−α12, and α13−α14) (Supplementary Figure S6). This structure also includes the PTS1-cargo binding region, which is a known feature of the Pex5 receptor TPR domain.

Details of the refinement statistics of the Eci1-Pex5 structure are described in Table S4. Three-dimensional visualization and analyses were performed using UCSF Chimera (Pettersen *et al*., 2004b) and PyMol (The PyMOL Molecular Graphics System, Version 2.0 Schrödinger, LLC. Available from: http://www.pymol.org/pymol). The coordinates of the Eci1 hexamer and the Eci1 (monomer)-Pex5 complex were deposited in the Protein Data Bank under the PDB codes 9FGZ and 9FH0, respectively. Their maps are available as EMD-50434 and EMD-50435.

## Data availability

The Eci1-Pex5 complex map and model have been deposited under accession codes EMD-50434 and 9FGZ, respectively. The Pex5 local refinement map and model have been deposited under accession codes EMD-50435 and 9FH0, respectively.

## Acknowledgments

LP is supported by the *Roy and Theo Caplan scholarship for promoting women students*. Work in the Schuldiner lab is supported by the ERC CoG OnTarget (864068). MS and EZ are supported by an Israeli Science Foundation grant 914/22. The robotic system of the Schuldiner lab was purchased through the kind support of the Blythe Brenden-Mann Foundation. MS is an Incumbent of the Dr. Gilbert Omenn and Martha Darling Professorial Chair in Molecular Genetics.

## Author contribution

LP, NE, AS, JJ, SA and YP performed the experiments; NE and OD analyzed the Cryo EM data, OD performed the structural analyses, SA, MS, YP and EZ supervised the work; OD, MS, YP and EZ conceptualized the work; LP, NE, OD, MS and EZ wrote the manuscript; All authors read and gave feedback on the manuscript.

---

**Figure S1.**
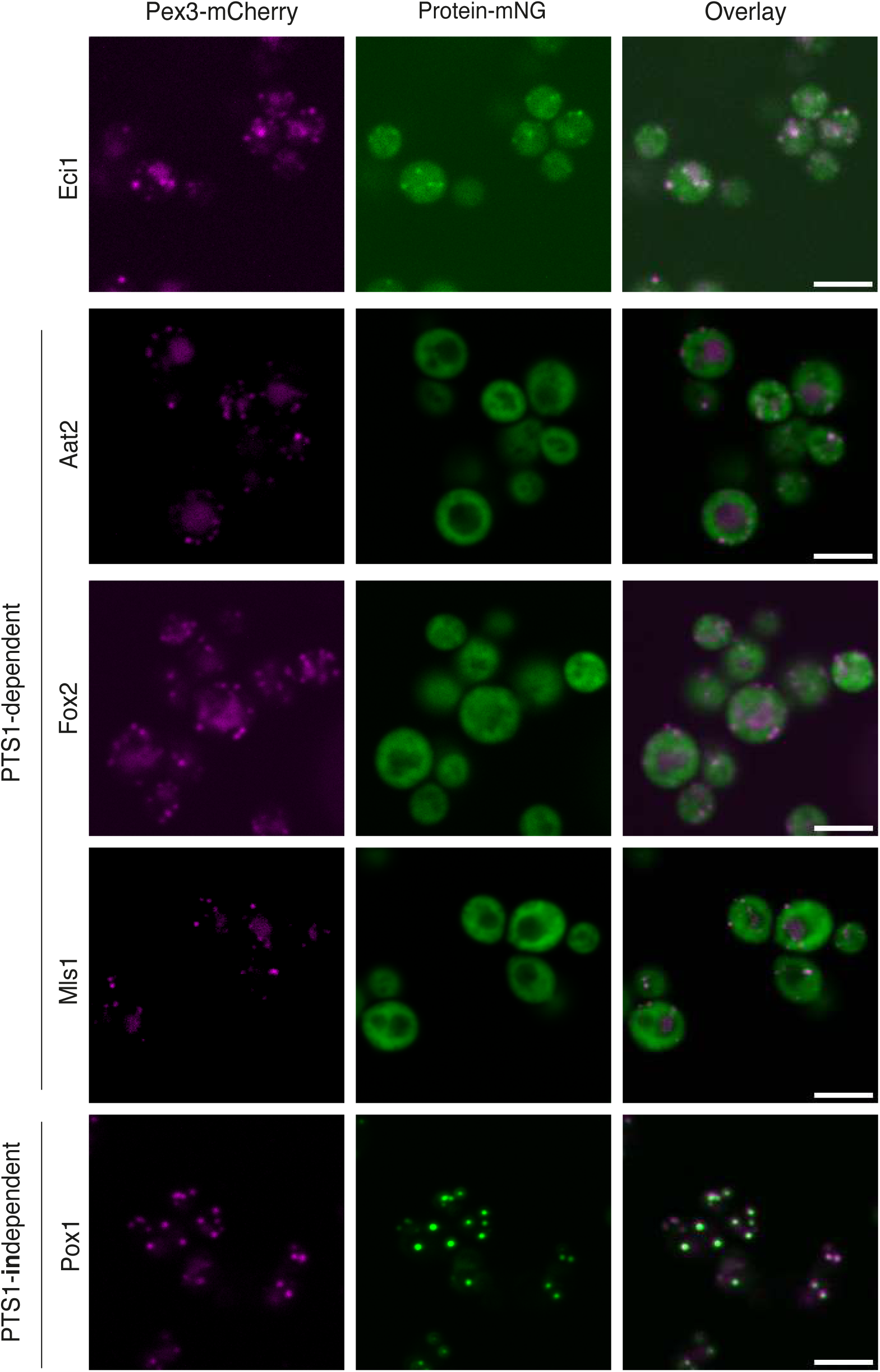
Proteins that are completely dependent on their PTS1 for targeting lose their peroxisomal localization upon masking their C terminus. Fluorescent microscopy images of peroxisomal proteins C’ tagged with mNG. Eci1-mNG maintains co-localization with the peroxisomal marker (Pex3-mCherry) suggesting an additional, PTS1 independent, mode of targeting. Proteins completely dependent on their PTS1 are no longer targeted to peroxisomes upon masking their PTS1 (i.e., Aat2, Fox2, and Mls1). Pox1, on the other hand, is targeted in a PTS1-independent manner, and hence, colocalizes with peroxisomes even when a fluorophore is masking its C terminus. Scale: 5 μm.

**Figure S2.**
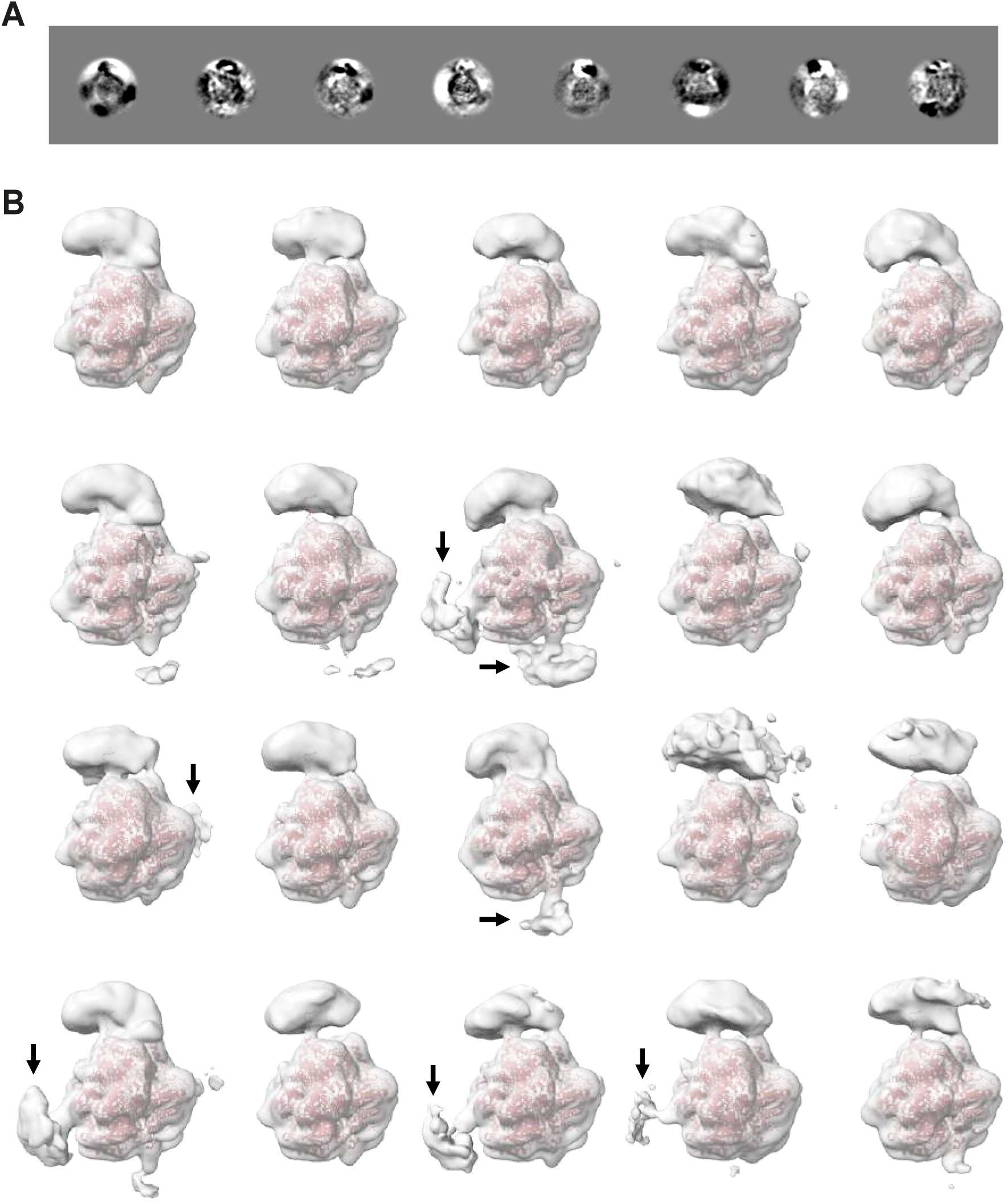
Analysis of Pex5-Eci1 stoichiometry in cryo-EM dataset. A) Eight eigenvectors (modes) calculated using 3D viability analysis, applying a spherical mask to include the Eci1 hexamer and surrounding Pex5 subunits. Contrast at the periphery of the mask indicates variability in Pex5 stoichiometry and/or conformation. B) Image dataset was separated into 20 clusters based on the above 8 eigenvectors and a single 3D map was calculated for each cluster. Eci1 coordinates were docked in for reference (red). The majority of Eci1 hexamers are bound by a single Pex5, while in some clusters more Pex5 are seen bound (pointed by arrows), all adopting a similar orientation relative to the hexamer.

**Figure S3.**
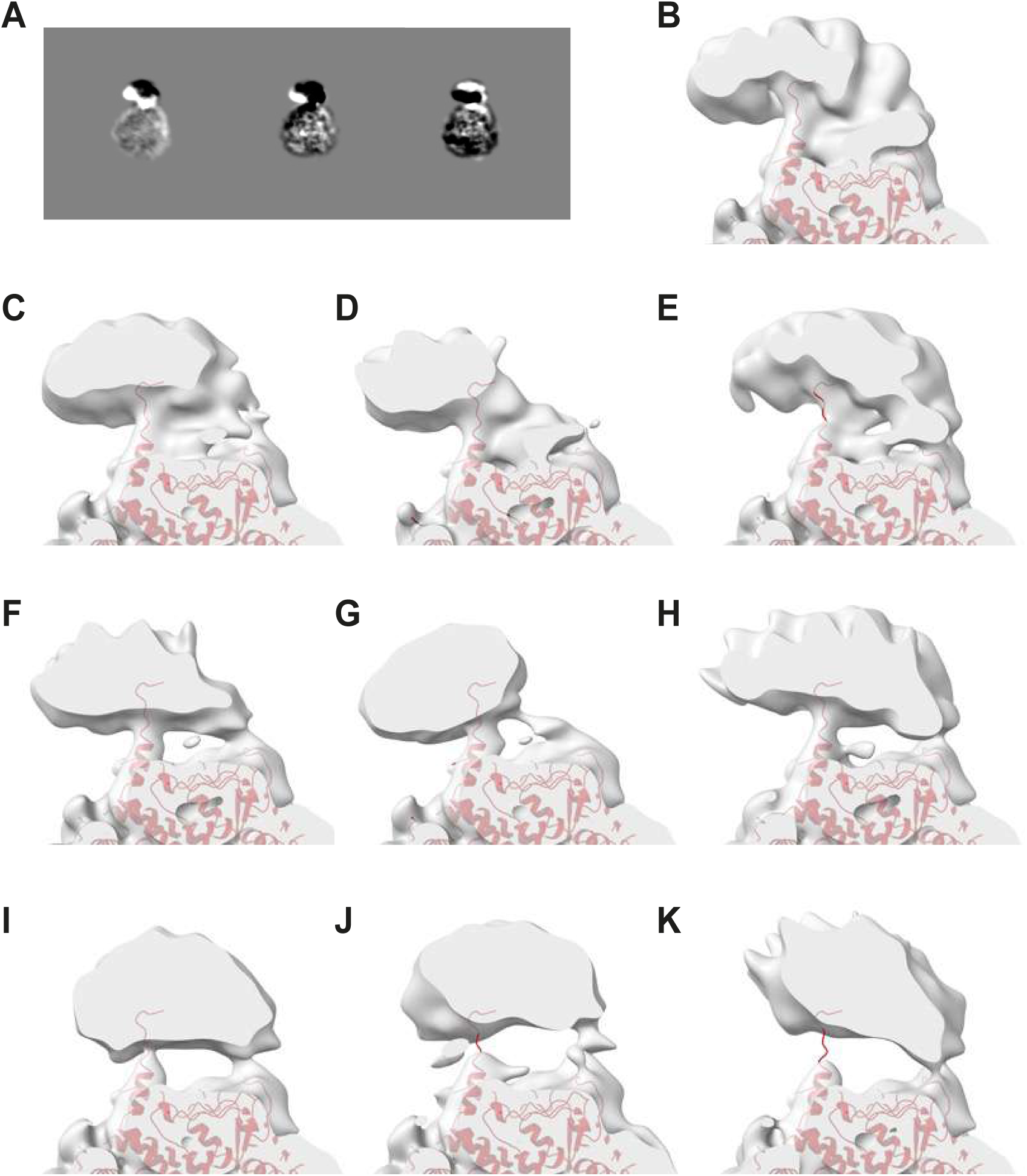
Pex5 conformational variability. A) Three eigenvectors (modes) calculated using 3D viability analysis, applying a mask to include the Eci1 hexamer and a single Pex5. The Pex5 region (top part) contains strong contrast, indicating significant conformational variability. B-K) Image data set was separated based on the above three eigenvectors into 10 clusters, and each cluster was subjected to 3D refinement. Shown are slabs through the refined maps, which were low-pass filtered to 10 Å. Eci1 coordinates were docked in for reference (red). Pex5 and the interface are best resolved in the map in panel B, where a continuous density appears between Eci1 and Pex5. This map was used for downstream processing. The interface and Pex5 are less well resolved in the other maps (C-K), although Pex5 remains anchored to Eci1 at two positions, one of them formed by the PTS1 signal peptide (F-K).

**Figure S4.**
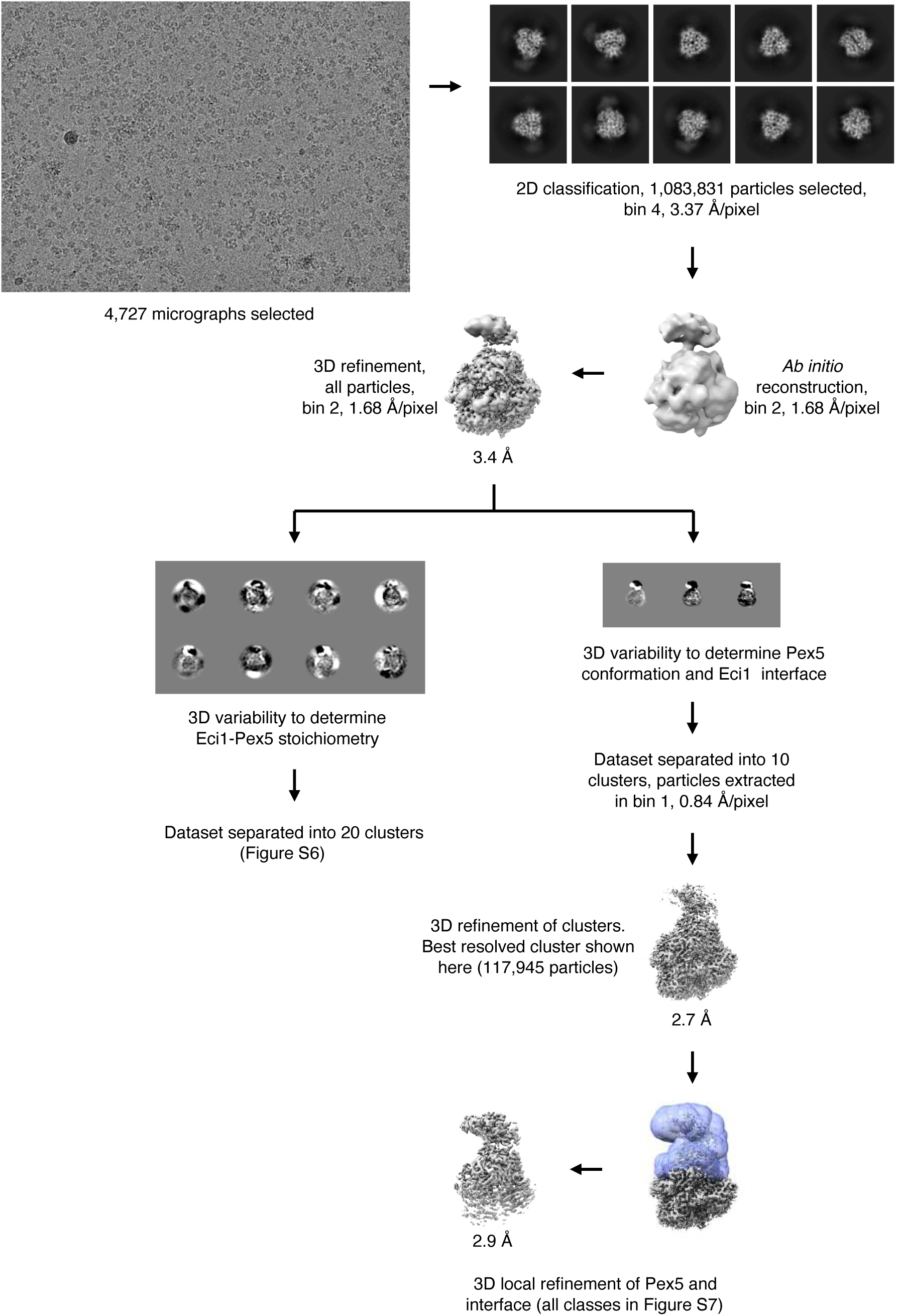
Single particle cryo-EM processing workflow. Shown is a graphical representation of the process as detailed in the methods section.

**Figure S5.**
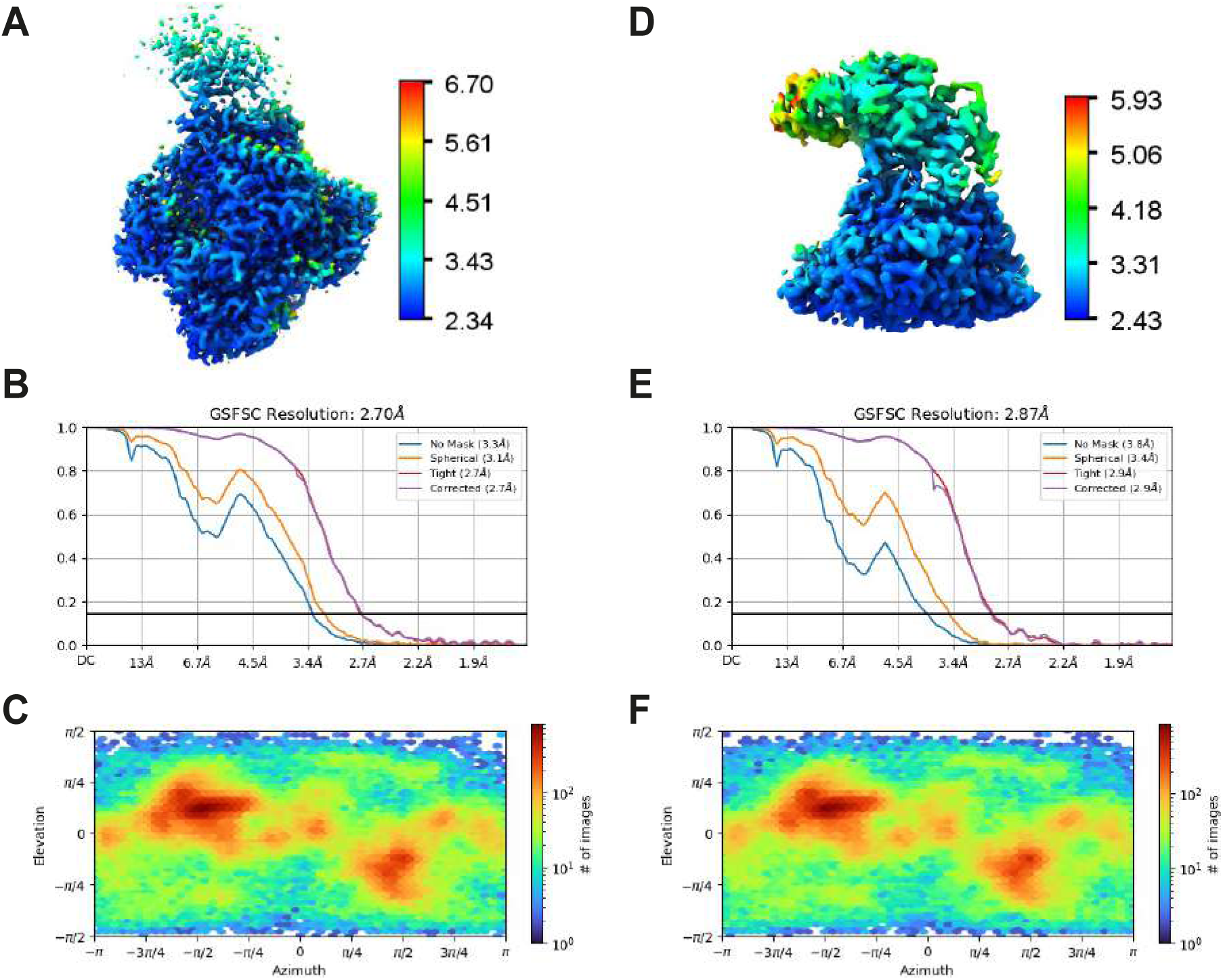
Refined cryo-EM maps used for model building. Plots for the full and local-refinement map are shown in panels A-C and D-F, respectively. A, D) 3D maps colored according to local resolution estimate. B, E) Fourier shell correlation (FSC) curves. C, F) Angular distribution plots.

**Figure S6.**
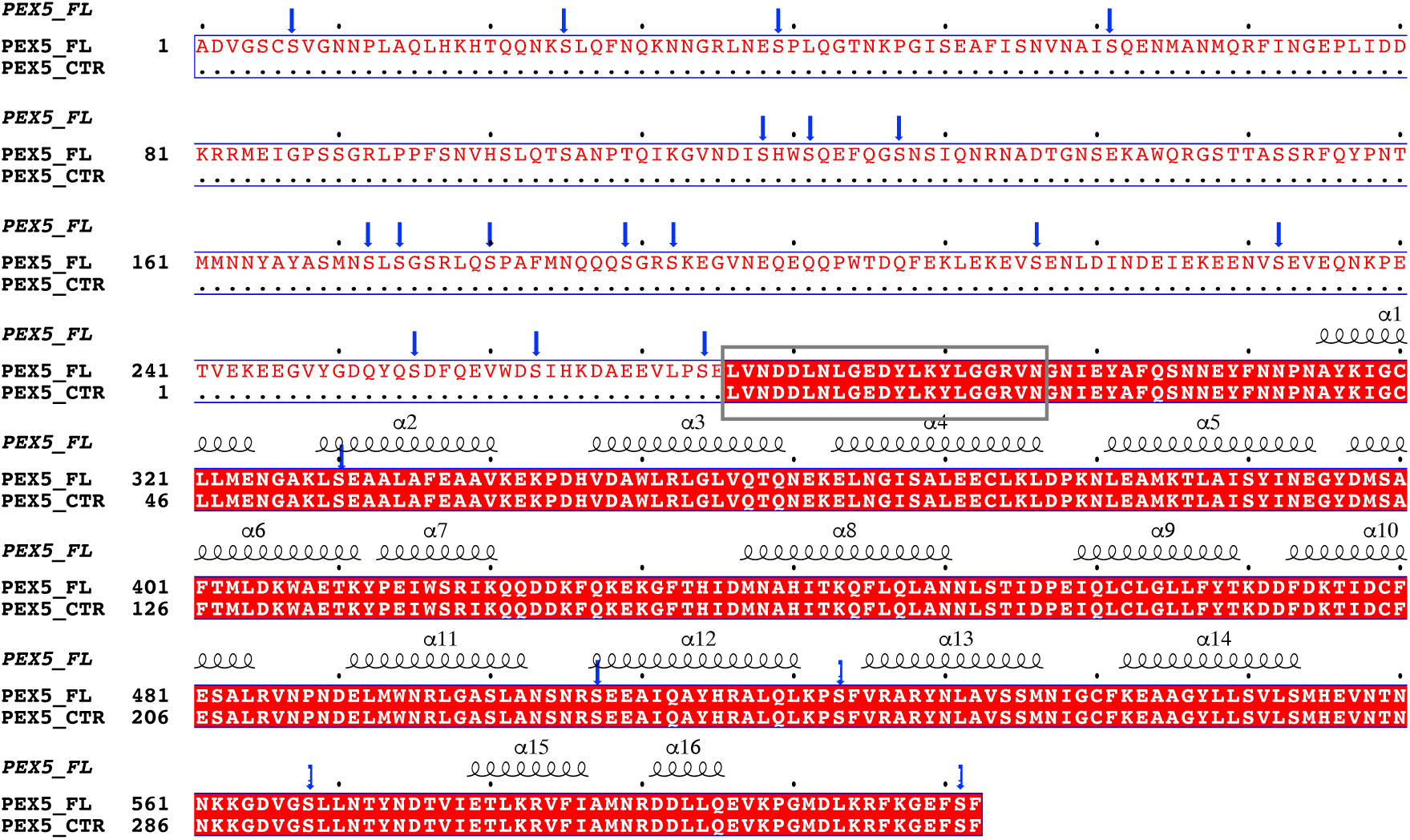
The resolved Pex5-Eci1 complex structure uncovers previously unseen residues of Pex5. The electron density-containing C-terminal domain of Pex5 (Pex5_CTR) was aligned with its full-length sequence (Pex5_FL), including the previously unobserved structure corresponding to the segment (^276^LVNDDLNLGEDYLKYLGGRVN^296^ depicted within the grey box) of Pex5. Secondary structure elements of the Cryo-EM structure of Pex5_CTR are labeled as follows: α-helices and 3_10_-helices (shown with the symbol “α”) are indicated by coils. Phosphorylated residues are highlighted with blue arrows. Multiple sequence alignment was performed using MultAlin (Corpet, 1988), and the figure was created using ESPript (Robert and Gouet, 2014)

**Figure S7.**
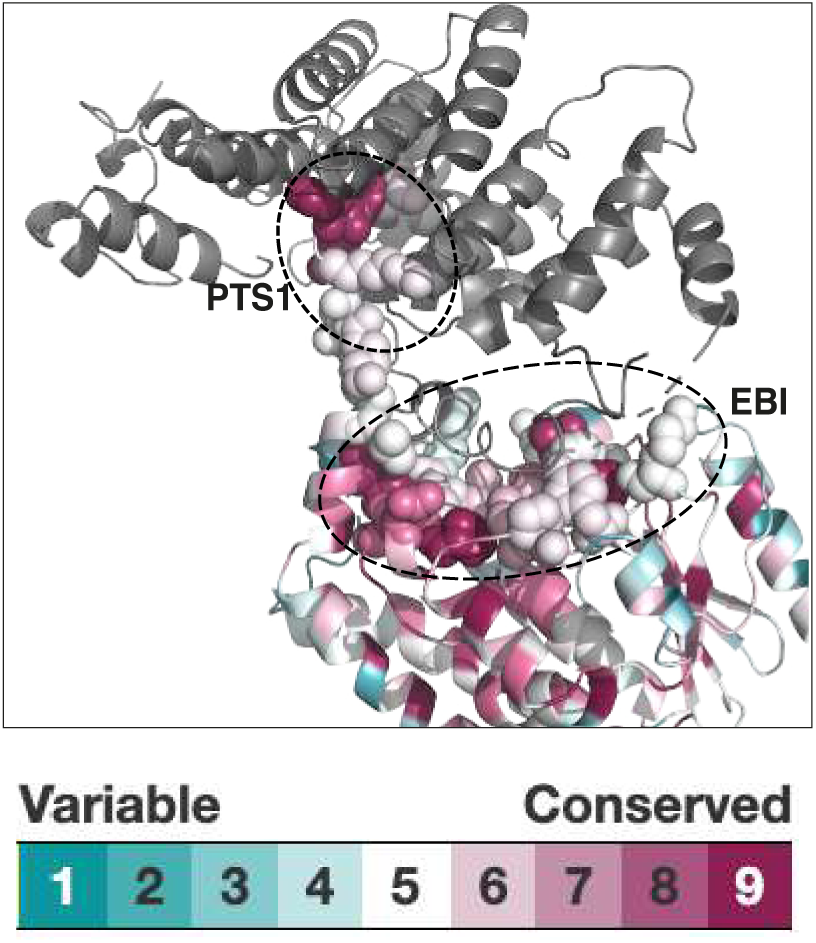
The residues of Eci1 involved in binding the PTS1 independent interface of Pex5 are conserved. Evolutionary conservation of the Cryo-EM resolved Eci1 structure is illustrated using the Consurf server (Ashkenazy *et al*., 2016), which estimates and visualizes evolutionary conservation in homologous proteins. Highly conserved amino acids are depicted in maroon, while the least conserved ones are shown in cyan. Notably, residues in Eci1 that interact with Pex5 in the two binding interfaces, the PTS1 binding interface and the EBI, are highly conserved. These interacting residues are represented as spheres.

**Figure S8.**
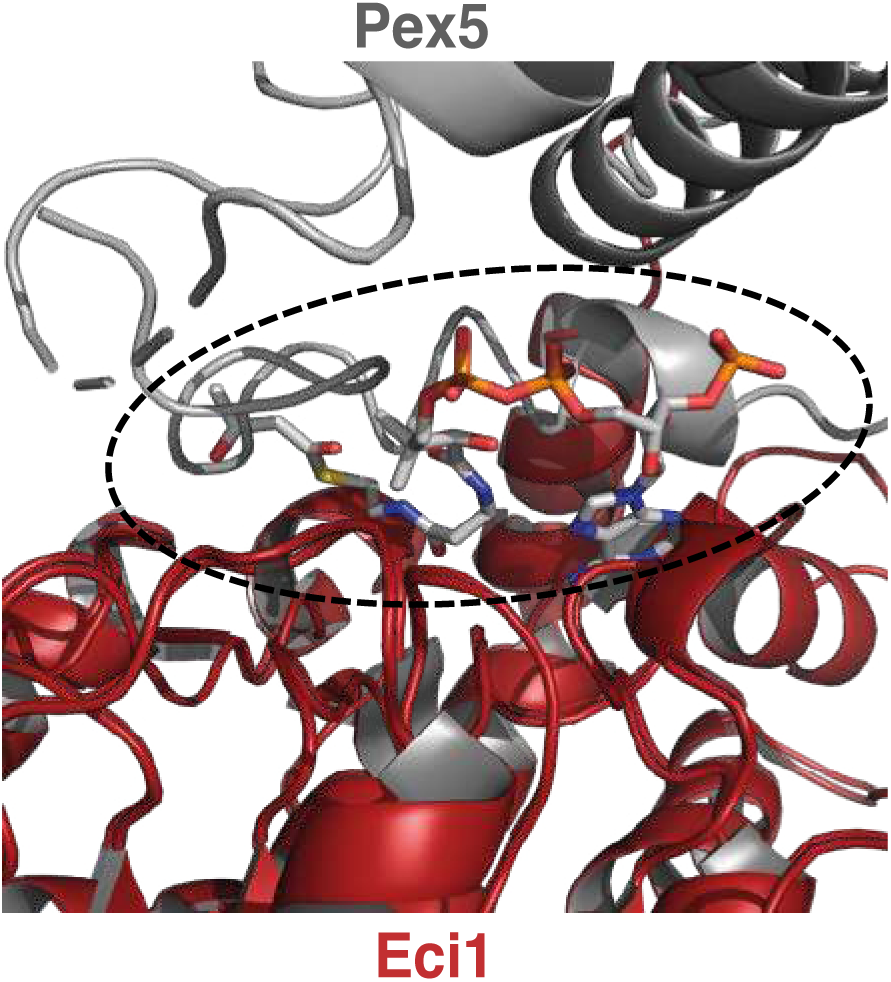
Eci1 residues interacting with Pex5 are also shared for substrate binding. The N-terminal segment of Pex5 (grey) occupies a position analogous to where CoA binds to Eci1 (red) in the Eci1-CoA complex (PDB entry 4ZDB), as indicated by the dashed black circle. CoA is represented as stick figures with carbon atoms colored grey, nitrogen atoms blue, oxygen atoms red, and phosphate atoms orange.

**Figure S9.**
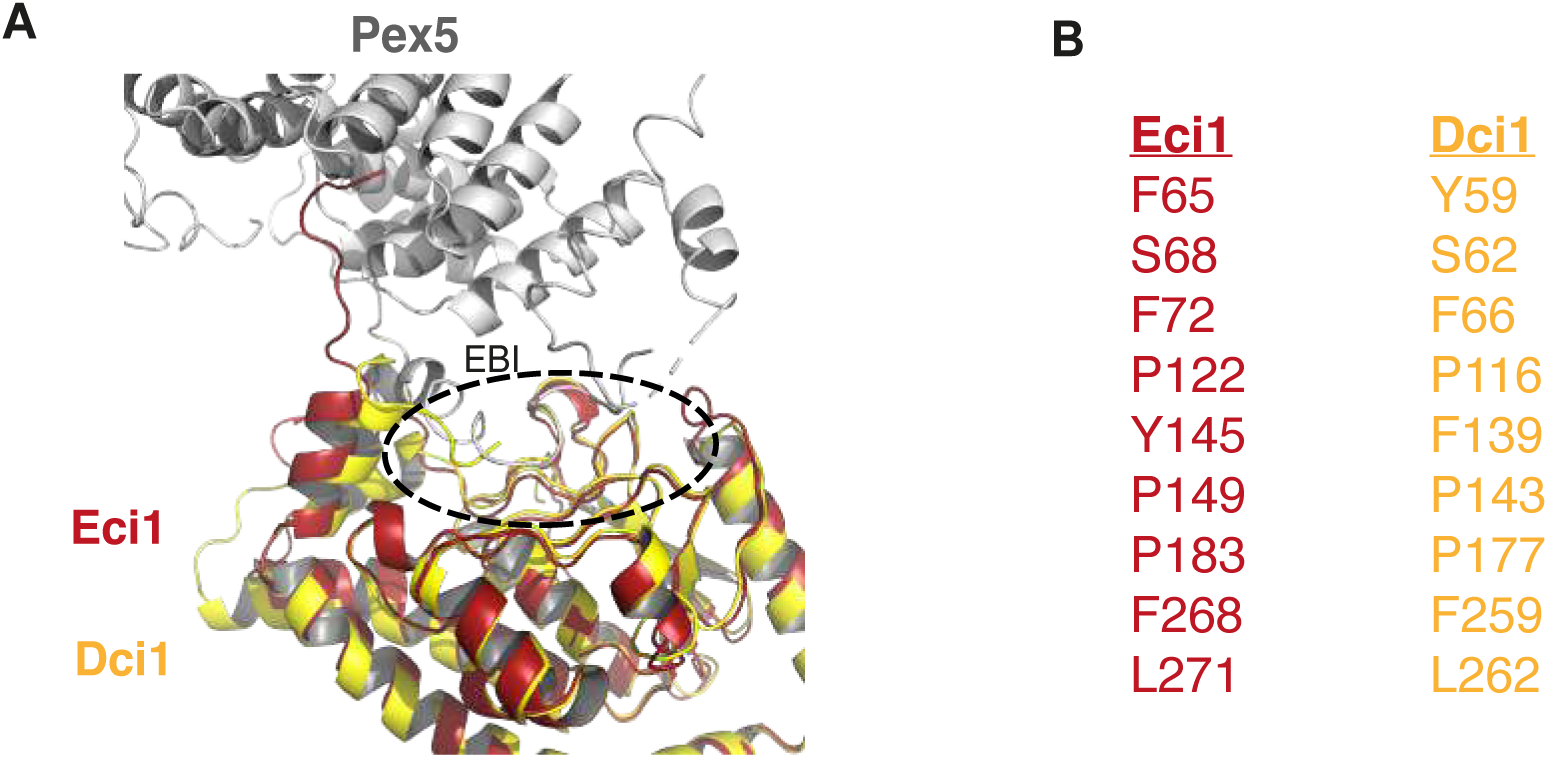
The Dci1 model exhibits alignment with Eci1 in residues involved in interaction with Pex5 via the novel EBI. A) The Dci1 predicted model (yellow) aligns well with the Cryo-EM structure of Eci1 (red). Remarkably, many residues of Eci1 that interact with Pex5 are highly conserved in its paralog Dci1.

## References

1. Abramson, J. et al. (2024) ‘Accurate structure prediction of biomolecular interactions with AlphaFold 3’, Nature [Preprint]. Available at: 10.1038/s41586-024-07487-w.

2. Adams, P.D. et al. (2010) ‘PHENIX: A comprehensive Python-based system for macromolecular structure solution’, Acta Crystallographica Section D: Biological Crystallography, 66(2), pp. 213–221. Available at: 10.1107/S0907444909052925.

3. Al-Hajaya, Y. et al. (2022) ‘Nuclear and peroxisomal targeting of catalase’, Plant Cell and Environment, 45(4), pp. 1096–1108. Available at: 10.1111/pce.14262.

4. Argyriou, C., D’Agostino, M.D. and Braverman, N. (2016) ‘Peroxisome biogenesis disorders’, Translational Science of Rare Diseases, 1(2), pp. 111–144. Available at: 10.3233/trd-160003.

5. Ashkenazy, H. et al. (2016) ‘ConSurf 2016: an improved methodology to estimate and visualize evolutionary conservation in macromolecules’, Nucleic Acids Research, 44(W1), pp. W344–W350. Available at: 10.1093/NAR/GKW408.

6. Aviram, N. and Schuldiner, M. (2017) ‘Targeting and translocation of proteins to the endoplasmic reticulum at a glance’, Journal of Cell Science, 130(24), pp. 4079– 4085. Available at: 10.1242/jcs.204396.

7. Bürgi, J. et al. (2023) ‘Asymmetric horseshoe-like assembly of peroxisomal Yeast Oxalyl-CoA synthetase’, Biological chemistry, 404, pp. 195–207. Available at: 10.1101/2022.08.30.505785.

8. Bykov, Y.S. et al. (2020) ‘Cytosolic Events in the Biogenesis of Mitochondrial Proteins’, Trends in Biochemical Sciences. Elsevier Ltd, pp. 650–667. Available at: 10.1016/j.tibs.2020.04.001.

9. Chen, V.B. et al. (2010) ‘MolProbity: All-atom structure validation for macromolecular crystallography’, Acta Crystallographica Section D: Biological Crystallography, 66(1), pp. 12–21. Available at: 10.1107/S0907444909042073.

10. Corpet, F. (1988) Nucleic Acids Research Multiple sequence alignment with hierarchical clustering.

11. Effelsberg, D. et al. (2016) ‘Pex9p is a new yeast peroxisomal import receptor for PTS1-containing proteins’, Journal of Cell Science, 129(21), pp. 4057–4066. Available at: 10.1242/jcs.195271.

12. Emsley, P. and Cowtan, K. (2004) ‘Coot: Model-building tools for molecular graphics’, Acta Crystallographica Section D: Biological Crystallography, 60(12 I), pp. 2126–2132. Available at: 10.1107/S0907444904019158.

13. Erdmann, R. and Blobel, G. (1995) Giant Peroxisomes in Oleic Acid-induced Saccharomyces cerevisiae lacking the Peroxisomal Membrane Protein Pmp27p.

14. Erijman, A. et al. (2011) ‘Transfer-PCR (TPCR): A highway for DNA cloning and protein engineering’, Journal of Structural Biology, 175(2), pp. 171–177. Available at: 10.1016/j.jsb.2011.04.005.

15. Fodor, K. et al. (2012) ‘Molecular requirements for peroxisomal targeting of alanine-glyoxylate aminotransferase as an essential determinant in primary hyperoxaluria type 1’, PLoS Biology, 10(4). Available at: 10.1371/journal.pbio.1001309.

16. Geisbrecht, B. V et al. (1998) Molecular Characterization of Saccharomyces cerevisiae 3 , 2-Enoyl-CoA Isomerase*. Available at: http://www.jbc.org.

17. Geisbrecht, B. V et al. (1999) Preliminary Characterization of Yor180Cp: Identification of a Novel Peroxisomal Protein of Saccharomyces cerevisiae Involved in Fatty Acid Metabolism. Available at: http://www.idealibrary.com.

18. Gould, S.J. et al. (1989) A Conserved Tripeptide Sorts Proteins to Peroxisomes.

19. Gurvitz, A. et al. (1998) Peroxisomal 3-cis-2-trans-Enoyl-CoA Isomerase Encoded by ECI1 Is Required for Growth of the Yeast Saccharomyces cerevisiae on Unsaturated Fatty Acids*. Available at: http://www.jbc.org.

20. Hanscho, M. et al. (2012) ‘Nutritional requirements of the BY series of Saccharomyces cerevisiae strains for optimum growth’, FEMS Yeast Research, 12(7), pp. 796–808. Available at: 10.1111/j.1567-1364.2012.00830.x.

21. Islinger, M. et al. (2018) ‘The peroxisome: an update on mysteries 2.0’, Histochemistry and Cell Biology. Springer Verlag, pp. 443–471. Available at: 10.1007/s00418-018-1722-5.

22. Jumper, J. et al. (2021) ‘Highly accurate protein structure prediction with AlphaFold’, Nature, 596(7873), pp. 583–589. Available at: 10.1038/s41586-021-03819-2.

23. Karpichev, I. V. and Small, G.M. (2000) ‘Evidence for a novel pathway for the targeting of a Saccharomyces cerevisiae peroxisomal protein belonging to the isomerase/hydratase famil’, Journal of cell science, 112(Pt3), pp. 533–544. Available at: 10.1242/jcs.113.3.533 (Accessed: 31 March 2024).

24. Kempiński, B. et al. (2020) ‘The Peroxisomal Targeting Signal 3 (PTS3) of the Budding Yeast Acyl-CoA Oxidase Is a Signal Patch’, Frontiers in Cell and Developmental Biology, 8. Available at: 10.3389/fcell.2020.00198.

25. Klaholz, B.P. (2019) ‘Deriving and refining atomic models in crystallography and cryo-EM: The latest Phenix tools to facilitate structure analysis’, Acta Crystallographica Section D: Structural Biology. International Union of Crystallography, pp. 878–881. Available at: 10.1107/S2059798319013391.

26. Klein, A.T.J. et al. (2002) ‘Saccharomyces cerevisiae acyl-CoA oxidase follows a novel, non-PTS1, import pathway into peroxisomes that is dependent on Pex5p’, Journal of Biological Chemistry, 277(28), pp. 25011–25019. Available at: 10.1074/jbc.M203254200.

27. Mastronarde, D.N. (2005) ‘Automated electron microscope tomography using robust prediction of specimen movements’, Journal of Structural Biology, 152(1), pp. 36–51. Available at: 10.1016/j.jsb.2005.07.007.

28. Mursula, A.M., Hiltunen, J.K. and Wierenga, R.K. (2004) ‘Structural studies on Δ3-Δ2-enoyl-CoA isomerase: The variable mode of assembly of the trimeric disks of the crotonase superfamily’, FEBS Letters, 557(1–3), pp. 81–87. Available at: 10.1016/S0014-5793(03)01450-9.

29. Onwukwe, G.U. et al. (2015) ‘Structures of yeast peroxisomal Δ3,Δ2-enoyl-CoA isomerase complexed with acyl-CoA substrate analogues: The importance of hydrogen-bond networks for the reactivity of the catalytic base and the oxyanion hole’, Acta Crystallographica Section D: Biological Crystallography. International Union of Crystallography, pp. 2178–2191. Available at: 10.1107/S139900471501559X.

30. Pettersen, E.F. et al. (2004a) ‘UCSF Chimera - A visualization system for exploratory research and analysis’, Journal of Computational Chemistry, 25(13), pp. 1605–1612. Available at: 10.1002/jcc.20084.

31. Pettersen, E.F. et al. (2004b) ‘UCSF Chimera - A visualization system for exploratory research and analysis’, Journal of Computational Chemistry, 25(13), pp. 1605–1612. Available at: 10.1002/jcc.20084.

32. Punjani, A. et al. (2017) ‘CryoSPARC: Algorithms for rapid unsupervised cryo-EM structure determination’, Nature Methods, 14(3), pp. 290–296. Available at: 10.1038/nmeth.4169.

33. Punjani, A. and Fleet, D.J. (2021) ‘3D variability analysis: Resolving continuous flexibility and discrete heterogeneity from single particle cryo-EM’, Journal of Structural Biology, 213(2). Available at: 10.1016/j.jsb.2021.107702.

34. Robert, X. and Gouet, P. (2014) ‘Deciphering key features in protein structures with the new ENDscript server’, Nucleic Acids Research, 42(W1). Available at: 10.1093/nar/gku316.

35. Rosenthal, M. et al. (2020) ‘Uncovering targeting priority to yeast peroxisomes using an in-cell competition assay’. Available at: 10.1073/pnas.1920078117/-/DCSupplemental.

36. Rymer, Ł. et al. (2018) ‘The budding yeast Pex5p receptor directs Fox2p and Cta1p into peroxisomes via its N-terminal region near the FxxxW domain’, Journal of Cell Science, 131(17). Available at: 10.1242/jcs.216986.

37. Senior, A.W. et al. (2020) ‘Improved protein structure prediction using potentials from deep learning’, Nature, 577(7792), pp. 706–710. Available at: 10.1038/s41586-019-1923-7.

38. Sonani, R.R. et al. (2023) ‘Noncanonical interactions and conformational dynamics in cargo-Pex5-Pex14 ternary complex for peroxisomal import’, bioRxiv, p. 2023.04.03.535445. Available at: 10.1101/2023.04.03.535445.

39. The PyMOL Molecular Graphics System, Version 2.0 Schrödinger, LLC. Available from: http://www.pymol.org/pymol (no date).

40. Unger, T. et al. (2010) ‘Applications of the Restriction Free (RF) cloning procedure for molecular manipulations and protein expression’, Journal of Structural Biology, 172(1), pp. 34–44. Available at: 10.1016/j.jsb.2010.06.016.

41. Walter, T. and Erdmann, R. (2019) ‘Current Advances in Protein Import into Peroxisomes’, *Protein Journal*. Springer Science and Business Media, LLC, pp. 351–362. Available at: 10.1007/s10930-019-09835-6.

42. Wanders, R.J. et al. (2023) THE PHYSIOLOGICAL FUNCTIONS OF HUMAN PEROXISOMES 2 3.

43. Waterham, H.R., Ferdinandusse, S. and Wanders, R.J.A. (2016) ‘Human disorders of peroxisome metabolism and biogenesis’, Biochimica et Biophysica Acta - Molecular Cell Research, 1863(5), pp. 922–933. Available at: 10.1016/j.bbamcr.2015.11.015.

44. Wei Xiao (2006) Yeast Protocols.

45. Williams, C.P. et al. (2011) ‘The peroxisomal targeting signal 1 in sterol carrier protein 2 is autonomous and essential for receptor recognition’, BMC Biochemistry, 12(1). Available at: 10.1186/1471-2091-12-12.

46. Yang, X., Purdue, P.E. and Lazarow, P.B. (2001) Eci1p uses a PTS1 to enter peroxisomes: either its own or that of a partner, Dci1p. Available at: www.urbanfischer.de/journals/ejcb.

47. Yifrach, E. et al. (2016) ‘Characterization of proteome dynamics during growth in oleate reveals a new peroxisome-targeting receptor’, Journal of Cell Science, 129(21), pp. 4067–4075. Available at: 10.1242/jcs.195255.

48. Yifrach, E. et al. (2022) ‘Systematic multi-level analysis of an organelle proteome reveals new peroxisomal functions’, Molecular Systems Biology, 18(9). Available at: 10.15252/msb.202211186.

49. Yifrach, E. et al. (2023) ‘Determining the targeting specificity of the selective peroxisomal targeting factor Pex9’, Biological Chemistry, 404(2–3), pp. 121–133. Available at: 10.1515/hsz-2022-0116.

50. Yofe, I. and Schuldiner, M. (2014) ‘Primers-4-Yeast: A comprehensive web tool for planning primers for Saccharomyces cerevisiae’, Yeast, 31(2), pp. 77–80. Available at: 10.1002/yea.2998.

51. Zahradník, J. et al. (2019) ‘Flexible regions govern promiscuous binding of IL-24 to receptors IL-20R1 and IL-22R1’, FEBS Journal, 286(19), pp. 3858–3873. Available at: 10.1111/febs.14945.

52. Zalckvar, E. and Schuldiner, M. (2022) ‘Beyond rare disorders: A new era for peroxisomal pathophysiology’, Molecular Cell. Cell Press, pp. 2228–2235. Available at: 10.1016/j.molcel.2022.05.028.

